# Robust diagnosis of infectious disease, autoimmunity and cancer from the paratope networks of adaptive immune receptors

**DOI:** 10.1101/2023.11.28.569125

**Authors:** Zichang Xu, Hendra S Ismanto, Dianita S Saputri, Soichiro Haruna, Guanqun Sun, Jan Wilamowski, Shunsuke Teraguchi, Ayan Sengupta, Songling Li, Daron M Standley

## Abstract

Liquid biopsies based on peripheral blood offer a minimally invasive alternative to solid tissue biopsies for the detection of diseases, primarily cancers. However, such tests currently consider only the serum component of blood, overlooking a potentially rich source of biomarkers: adaptive immune receptors (AIRs) expressed on circulating B and T cells. Machine learning-based classifiers trained on AIRs have been reported to accurately identify not only cancers, but also autoimmune and infectious diseases as well. However, when using the conventional “clonotype cluster” representation of AIRs, donors within a disease or healthy cohort exhibit vastly different features, limiting the generalizability of these classifiers. This paper addresses the challenge of classifying specific diseases from circulating B or T cells by developing a novel representation of AIRs based on similarity networks constructed from their antigen-binding regions (paratopes). Features based on this novel representation, paratope cluster occupancies (PCOs), significantly improved disease classification performance for infectious disease, autoimmunity and cancer. Under identical methodological conditions, classifiers trained on PCOs achieved a mean ROC AUC of 0.893 when applied to new donors, compared to clonotype cluster-based classifiers (0.714) or the best-performing published classifier (0.777). Surprisingly, for cancer patients, we observed that some of the AIRs that were important for classification were significantly more abundant in healthy controls than in individuals with disease. These “healthy-biased” AIRs were predicted to target known cancer-associated antigens at dramatically higher rates than healthy AIRs as a whole (Z scores > 75), suggesting the existence of an overlooked reservoir of cancer-targeting immune cells that are diagnostic and identifiable from a routine blood test. Consequently, PCOs not only enhance classification of a broad range of diseases but also identify immune cells with therapeutic potential.

## Introduction

Liquid biopsies that extract circulating tumor DNA, extracellular vesicles, or circulating tumor cells from peripheral blood offer a range of advantages over traditional methods of medical diagnosis^1^. Compared to solid tissue biopsies, these test are minimally invasive, which facilitates routine monitoring, which offers the promise of disease detection before clinical symptoms manifest^2^. More broadly, such approaches have the potential to empower individuals to manage their own health and thus to reduce the cost of and accessibility to healthcare. Furthermore, liquid biopsies may provide molecular profiles of heterogeneous diseases, which can inform personalized treatment interventions^3^. Finally, technological improvement of next-generation sequencing (NGS) and machine learning (ML) are expected to further accelerate improvements in the sensitivity and specificity of these tests^4^.

Despite their great promise, current blood-based liquid biopsies have limitations as well. The sensitivity of circulating tumor DNA detection can be low, especially in early-stage cancers or diseases with low tumor burden^5^. Another shortcoming is the exclusive focus on serum. Blood is a rich source of immune cells, which play a direct role in responses to many diseases. Incorporating the sequences of adaptive immune receptors (AIRs) would greatly enrich the information content of liquid biopsies, potentially improving diagnostic sensitivity^6^. As AIR sequencing and computational analysis technologies have continued to improve, application of AIR data has grown from basic research to application to biomarkers for disease and for guiding immunotherapies^7^. Taken together, expanding current liquid biopsies to include AIR sequence information warrants further exploration.

Each adaptive immune cell expresses a unique AIR, whose coding sequence is generated by rearrangement of germline V D and J genes^8^ (**Fig. S1A**). The number of possible combinations far exceeds the number of B or T cells in any one individual^9, 10^. The resulting extraordinary diversity allows adaptive immune cells to engage with and remember nearly any disease-associated antigen. Upon antigen engagement, adaptive immune cells proliferate, resulting in dramatic differences in the levels of specific AIR clones in peripheral blood. For use as a liquid biopsy, the diverse AIR signals in a given donor must be formulated as a single feature vector that can be compared to those of other donors. In recent years, statistics- and ML-based approaches have been actively explored in order to construct such features in order to classify donors according to their disease status^11–20^.

The main obstacle to the use of AIRs features of disease status is their diversity. The pairwise sharing of AIRs between different donors from typical blood samples ranges from 1-6% and decreases rapidly with an increase in the number of donors^21, 22^. This “donor sharing problem” has severely hindered the use of AIRs as traditional biomarkers as it prevents ML classifiers from being able to identify general features associated with particular disease. The extent of donor sharing depends, however, on the way of constructing the AIR features. The traditional representation of an AIR is as “clonotype”: its V and J gene names, along with its CDR3 amino acid sequence. A clonotype is a useful qualitative nomenclature because it describes the receptor’s gene rearrangement history. However, because clonotypes mix categorical (gene names) and continuous (CDR3 amino acid sequence) variables, it is not ideal for quantifying the similarities or differences between AIRs.

An alternative approach is to represent AIRs by a single amino acid sequence containing the residues near the antigen binding interface, also known as the “paratope”. Paratopes bring together three distinct segments called complementarity determining regions (CDR1, CDR2, add CDR3) (**Fig. S1B**). Most physical contacts between AIRs and antigens occur within or near the CDRs (**Fig. S1C**). By concatenating the three complementarity regions^23^ or by predicting the paratope^24^ a single paratope sequence can be constructed that is readily handled by standard protein sequence analysis methods for alignment, clustering, or searching^25, 26^. This approach has thus been used for grouping BCRs that target a common antigen^23, 24, 27, 28^ or TCRs that target a given peptide-MHC complex^17, 29^. In the context of disease diagnosis, an important strength of the paratope representation is that AIRs form extended networks that join together different clonotypes and, importantly, different donors.

## Results

### Paratope adjacencies connect diDerent clonotypes and donors

The impact of the paratope representation is best seen from an example. For this purpose, we constructed clonotype (**Fig. 1A**) and paratope (**Fig. 1B**) networks using the BCRs of ten COVID-19 from a previous study^30^. Notably, none of the clonotype networks connected different donors, consistent with the previously reported low sharing of BCR clones^9, 22^. The frequency of cluster sizes was systematically lower for clonotypes, than for paratopes, which formed a number of very large networks (**Fig. 1C**). Importantly, the larger paratope networks often contained many different clonotypes and connected many donors (**Fig. 1D**). In this context, we refer to the AIRs that are connected in the paratope networks as “adjacent”. These observations on clonotype and paratope networks form the basis of the features introduced here. The key observation is that AIRS can be “paratope adjacent” even if they come from different donors. Therefore, features based on such adjacency are likely to have elements that are shared among different donors. This is essential to group donors belonging to a common disease class.

**Figure 1.**
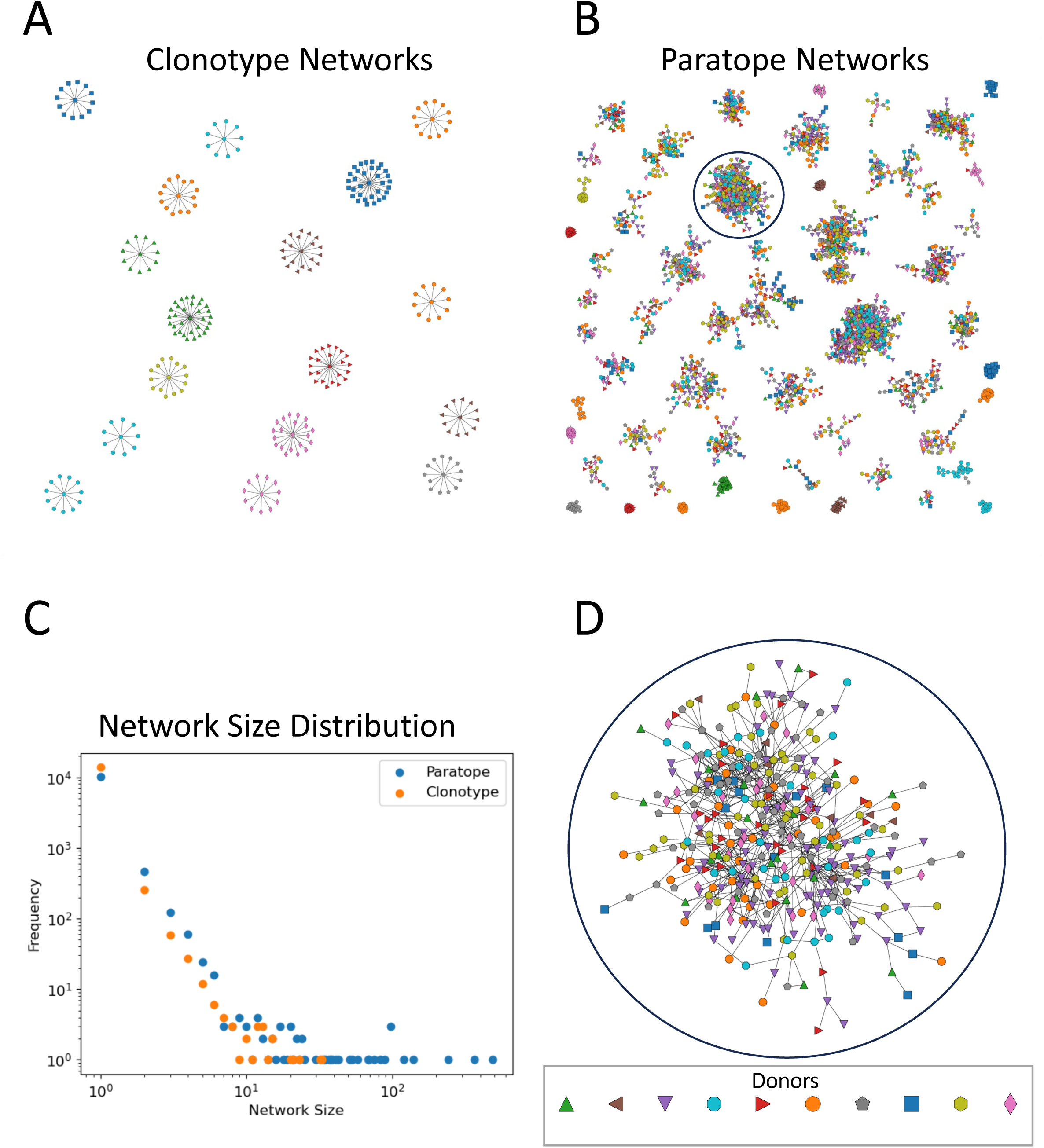
Paratope networks enable donor sharing. BCR heavy chain networks were constructed for 10 COVID-19 patients, where nodes are colored by donor and edges represent BCR sharing the same clonotype or similar paratope. Only networks larger than 10 are shown for simplicity. **A,** Clonotype networks are typically small and do not connect different donors. **B,** Paratope networks are much larger and generally connect many donors. **C,** The network sizes and frequencies were distinct even on a log scale. **D,** A close-up view of one of the larger paratope networks (circled in B), whch is made up of many different clonotypes, includes all 10 donors.

### Feature engineering

We describe the details of our approach in the *Methods* section. Briefly, our goal is to derive a single feature vector for each donor. We assume each AIR belongs to a cluster. Such clusters can be defined in terms of clonotypes or paratopes. An individual donor can be represented by the frequency of clusters. We denote such features as “cluster frequencies” (CFs) (**Fig. S2A**). In this way, CFs consist of “clonotype cluster frequencies” (CCFs) or “paratope cluster frequencies” (PCFs)), depending on whether we use clonotype or paratope clustering. We next extend the CFs by considering the networks of paratopes (**Fig. 1B**). The networks are represented mathematically by an adjacency matrix 𝐴 of pairwise paratope edges (**Fig. S2B**). Importantly, a given AIR can have edges with AIRs in the same cluster as well to AIRs belonging to different clusters. We denote the frequency of edges to all clusters as “occupancies” to emphasize the notion that a given AIR can “occupy” multiple clusters. Cluster occupancies (COs) consist of “clonotype cluster occupancies” (CCOs) or “paratope cluster occupancies” (PCOs), depending on whether we use clonotype or paratope clustering (**Fig. S2C**). Because all features are indexed by a specific cluster, the feature importance, as calculated by XGBoost, can be used to identify biologically important AIRs.

### AIR data downsizing

Because AIR frequencies follow a long-tailed distribution (**Fig. S1D**), cluster frequencies also have a long-tailed distribution, which results in very long feature vectors. We thus require a way to drop clusters that are not populated by many donors. We introduce a parameter 𝑑_min_ that sets a threshold for the required donor diversity. By discarding features with low donor diversity, we can reduce the sparsity of the features, reduce the dimensionality of the feature vectors, and reduce the number of AIRs in the paratope adjacency matrix. The parameter 𝑑_min_ is thus tuned in the training process, as described in the *Results* section.

To evaluate the classifiers for a wide range of diseases, we collected AIR sequences for 6 diseases covering 3 general categories (**Table S1**): infectious disease (COVID-19, HIV), autoimmune disease (autoimmune hepatitis, type 1 diabetes), and cancer (colorectal cancer, non-small cell lung cancer). Wherever possible, we utilized disease and healthy control data from the same study to minimize batch effects (i.e., any bias other than that of the disease of interest) or used different studies for training and testing.

### Classifier training

We randomly split the donors into training (70%) and test (30%) groups. For the classifiers described here, we separately selected the optimal hyperparameter 𝑑_min_ by leave-one-out cross validation (LOOCV) using the training donors. Two previously published methods, DeepRC^20^ and immuneML^19^, were identically trained, as described in the *Methods* section. We then evaluated all the trained classifiers on the test set (**Fig. S2C**). Aside from the feature generation steps (CCF, CCO, PCF, and PCO), all steps in the pipeline were identical for all of the newly developed classifiers. In the following sections, we discuss two representative diseases—COVID-19 (using single cell BCR sequence data) and NSCLC (using bulk TCR sequence data)—in depth. Our findings for the remaining diseases are described in **Figs S3-6 and the Suppl text**. XXX

### Classification of COVID-19 patients and healthy donors from BCR data

COVID-19 is caused by severe acute respiratory syndrome coronavirus 2 (SARS-CoV-2), which initially targets the epithelial tissue of the nasal cavity and spreads to the upper respiratory tract, lung, and other organs^31^. Diagnosis is usually confirmed by polymerase chain reaction (PCR), which can detect the presence of viral nucleic acids^32^. To evaluate the performance of a theoretical AIR-based diagnostic system, we utilized BCR heavy chain data from a single-cell sequencing study of peripheral blood mononuclear cells (PBMCs) from 44 COVID-19 patients (106,640 BCRs) and 58 healthy donors (174,139 BCRs), all of whom were unvaccinated^30^. After splitting the data randomly, 71 donor datasets were used for training, and 31 were used for testing. The sparsity of the features decreases roughly linearly with 𝑑_min_, while the LOOCV target function reached a maximum at a 𝑑_min_value of 0.7 (**Fig. 2A**). Using this value of 𝑑_min_,the classifier trained on the PCO features achieved an area under the receiver operating characteristic curve (ROC AUC) of 0.896 for test data, which was close to the LOOCV result on training data (0.946), indicating that the classifier was not overfitted and generalized well to new donors (**Fig. 2B**). When compared with the other new classifiers, only the model trained on CCF features was unable to classify the test donors (**Fig. 2C)**, presumably due to the poor sharing of clonotypes across donors, as described above. The performance of the previously published classifiers ranged from 0.407 to 0.838, highlighting the importance of the technology in achieving robust results. The precision‒recall (PR) AUCs indicate that the PCO-based classifier achieved the highest value (0.932) among all the trained models (**Fig. 2D)**. The above results demonstrate the improved generalizability of the PCO-based classifier, in particular in comparison to the CCF-based classifier. XXX

**Figure 2.**
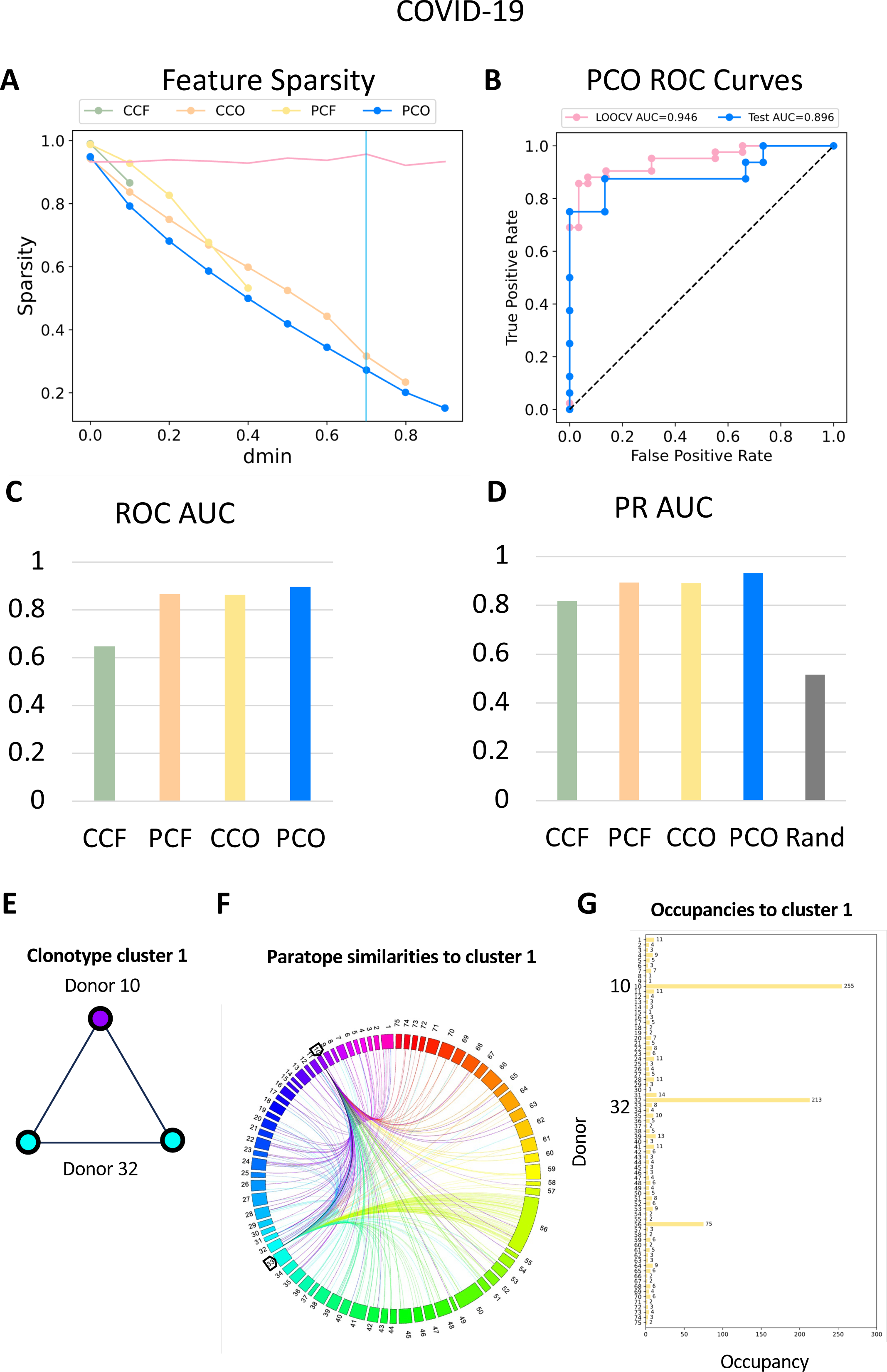
Assessment of COVID-19 diagnosis based on BCRs. **A,** The sparsity of each of the four feature matrices as a function of 𝑑_min_ . The vertical cyan line shows the value from LOOCV hyperparameter optimization, and the pink line indicates the target function (mean of training ROC AUC and PR AUC). **B,** ROC curves for the LOOCV and test predictions using the PCO classifier. **C,** ROC AUCs for all four classifiers in the test set. **D,** PR AUCs for all four classifiers in the test set; the performance of a random classifier is indicated as a gray bar. **E,** Impact of occupancy on important features. The clonotype cluster corresponding to the most important feature in the COVID-19 CCO training (cluster 1) has three members representing two donors (donor 10 and donor 32). The small size and low sharing of cluster 1 result in a sparse CCF feature. **F,** AIRs with similar paratopes from other clonotype clusters are visualized as a chord diagram, where each color denotes a different donor. The original two donors are colored purple and cyan, as in the clonotype cluster. **G,** A high degree of paratope-level similarity across multiple BCRs from multiple donors results in a dense CCO feature vector.

**Figure 3.**
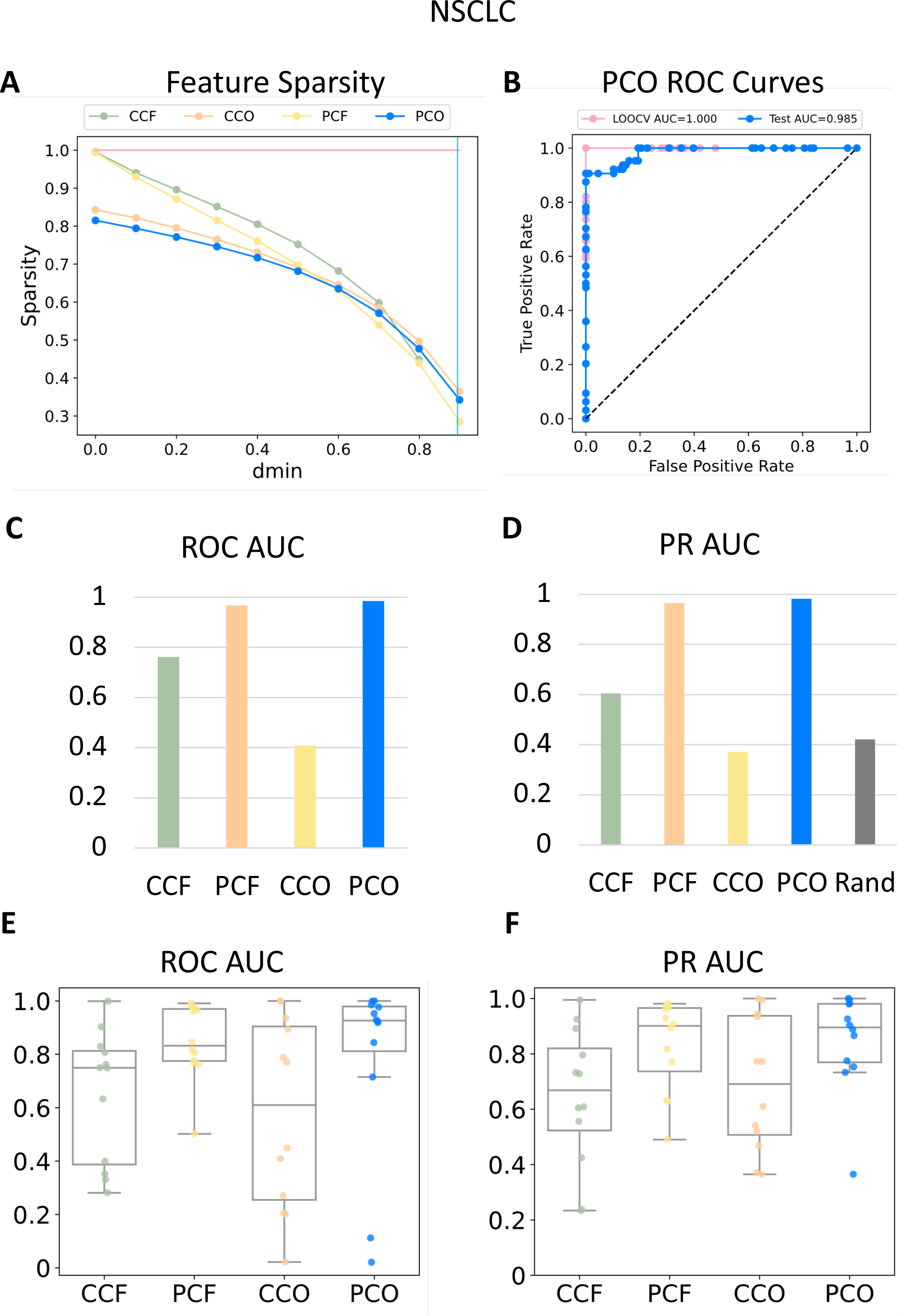
Assessment of NSCLC diagnosis based on TCRs. **A**, The sparsity of each of the four feature matrices as a function of 𝑑_min_, with the vertical cyan line showing the value from LOOCV hyperparameter optimization and the pink line indicating the target function (mean of training ROC AUC and PR AUC). **B,** ROC curves for the LOOCV and test predictions using the PCO classifier. **C,** ROC AUCs for all four classifiers in the test set. **D,** PR AUCs for all four classifiers in the test set; the AUC of a random classifier is indicated as a gray bar. **E,** Summary of 12 ROC AUC values in the test sets of NSCLC using different combinations of studies for training and testing. **F,** PR AUC values in the same 12 test sets.

The stark difference in performance between the CCF-based classifiers and the remaining types classifiers suggests that the computation of paratope occupancies had a critical impact on differentiating healthy individuals from those with COVID-19. To understand this effect, we examined differences between CCF and CCO features, the latter of which utilized the same clonotype clusters as the former but were transformed to occupancies using paratope similarities. To this end, we first identified the feature with the highest CCO importance. We subsequently examined the clonotype cluster (Cluster 1) corresponding to this feature. This cluster consisted of only three AIRs from 2 of 104 possible donors: donor 10 and donor 32 (**Fig. 2E)**. By examining the CCO calculations, we found that AIRs from many donors shared paratope-level similarity with one or more of the three members of cluster 1 (**Fig. 2F**). Thus, while the CCF feature corresponding to this cluster was zero for all but the two donors (10 and 32), the corresponding CCO feature was nonzero for many donors (898 AIRs from 75 of 104 donors) (**Fig. 2G**). This example highlights the fact that an AIR that belongs to a given clonotype cluster can also have significant paratope similarity to AIRs in different clusters and that including such paratope-level similarities was therefore critical to the good performance of the CCO- and PCO-based COVID-19 classifiers.

### Classification of Non-small cell lung cancer patients from healthy donors using TCR data

Non-small cell lung cancer (NSCLC) is the most common type of lung cancer; it consist primarily of squamous cell carcinoma, large cell carcinoma, and adenocarcinoma, but several rarer subtypes have also been identified^33^. Diagnosis traditionally relies upon a wide variety of evidence, including symptoms (e.g., persistent cough), imaging data, and tissue biopsy; however, diagnoses based on blood-based biomarkers (*EGFR, HER2, BRAF, KRAS, MET etc*)^34^ have recently become more common in the application of liquid biopsy in NSCLC^35^. Like in many other cancers, in NSCLC, disease-specific T cells often infiltrate tumors, but whether these T cells also circulate in the blood remains poorly understood. A TCR-based diagnosis would be beneficial as a less invasive approach than biopsy. Unlike for the other diseases, we were unable to find a single study that included PBMCs from both NSCLC patients and healthy controls. Therefore, we constructed a diverse dataset from seven studies: three with data from healthy individuals^36–38^ and two with data from individuals with NSCLC^39, 40^ to serve as the training set and one each with data from healthy individuals^41^ and from individuals with NSCLC^42^ to serve as the testing set. Across the studies, a total of 204 NSCLC donors (6,734,867 TCRs) and 294 healthy donors (4,120,597 TCRs) were ultimately included. After the data were split, 344 donors were used for classifier training, and 152 were retained for testing. **Fig. 3A** shows the sparsity of the resulting features, indicating that the occupancy-based features (CCO, PCO) were approximately 20% less sparse than the frequency-based features at small values of 𝑑_min_ and converged at larger values, while the LOOCV target function remained perfect at all values. **Fig. 3B** shows that the PCO-based classifier trained using the highest 𝑑_min_(0.9) achieved an ROC AUC of 0.985 for the test donors. **Figs. 3C-D** indicate that, among the new classifiers, only the those trained on occupancy-based features (CCO, PCO) performed well on test donors and that their performance was close to perfect (ROC AUC 0.985; PR AUC 0.982). The ROC AUCs of the previously published methods ranged from 0.317 to 0.895. Taken together, PCO-based classifiers demonstrated the potential for nearly perfect diagnosis on a large cohort of cancer patients and healthy controls. The less than perfect performance of the next-best classifier--0.895 by immuneML(RF)—indicated that merely increasing the size of the training data was not sufficient for robust classification.

### Assessment across infectious disease, autoimmunity and cancer

In addition to COVID-19 and NSCLC, we extended our benchmark to include an additional infectious disease, Human immunodeficiency virus (HIV); autoimmunity (Autoimmune hepatitis (AIH), Type 1 Diabetes (T1D); and, an additional cancer, Colorectal cancer (CRC) (**Figs. S3-4**). When we assessed all classifiers on all six independent test donor sets, we found that the PCO-based classifier exhibited the best overall performance (**Table 1**). The PCO-based classifier achieved the greatest ROC AUC among all the classifiers for all six diseases except for CRC, the disease with the fewest patients (20), and for which the difference (0.850 vs 0.821) was rather small. The mean ROC AUC of the PCO-based classifier (0.893) was substantially greater than that of the CCF-based classifier (0.714) and the previously published methods (0.537-0.777). Three of the PCO ROC AUCs—those for HIV (0.985), autoimmune hepatitis (AIH, 0.947) and NSCLC (0.985)—were close to perfect. The remaining three diseases—COVID-19 (0.896), type 1 diabetes (T1D, 0.725), and CRC (0.821)—were represented by fewer disease donors, suggesting that at least 50 disease donors may be needed for robust classifier performance. In contrast, the CCF-based classier ROC AUCs exceeded 0.9 for only one disease (HIV), for which the sequencing depth was much greater than that of the remaining diseases and performed no better than chance for T1D. It is noteworthy that, removal of the paratope features reduced performance (CCF, 0.714) to a value similar to that of immuneML(RF) (0.777), suggesting that the improvement was likely due to this innovation. The above results demonstrate that PCO-based classifiers can successfully distinguish disease patients from healthy donors in infectious disease, autoimmunity and cancer.

**Figure 4.**
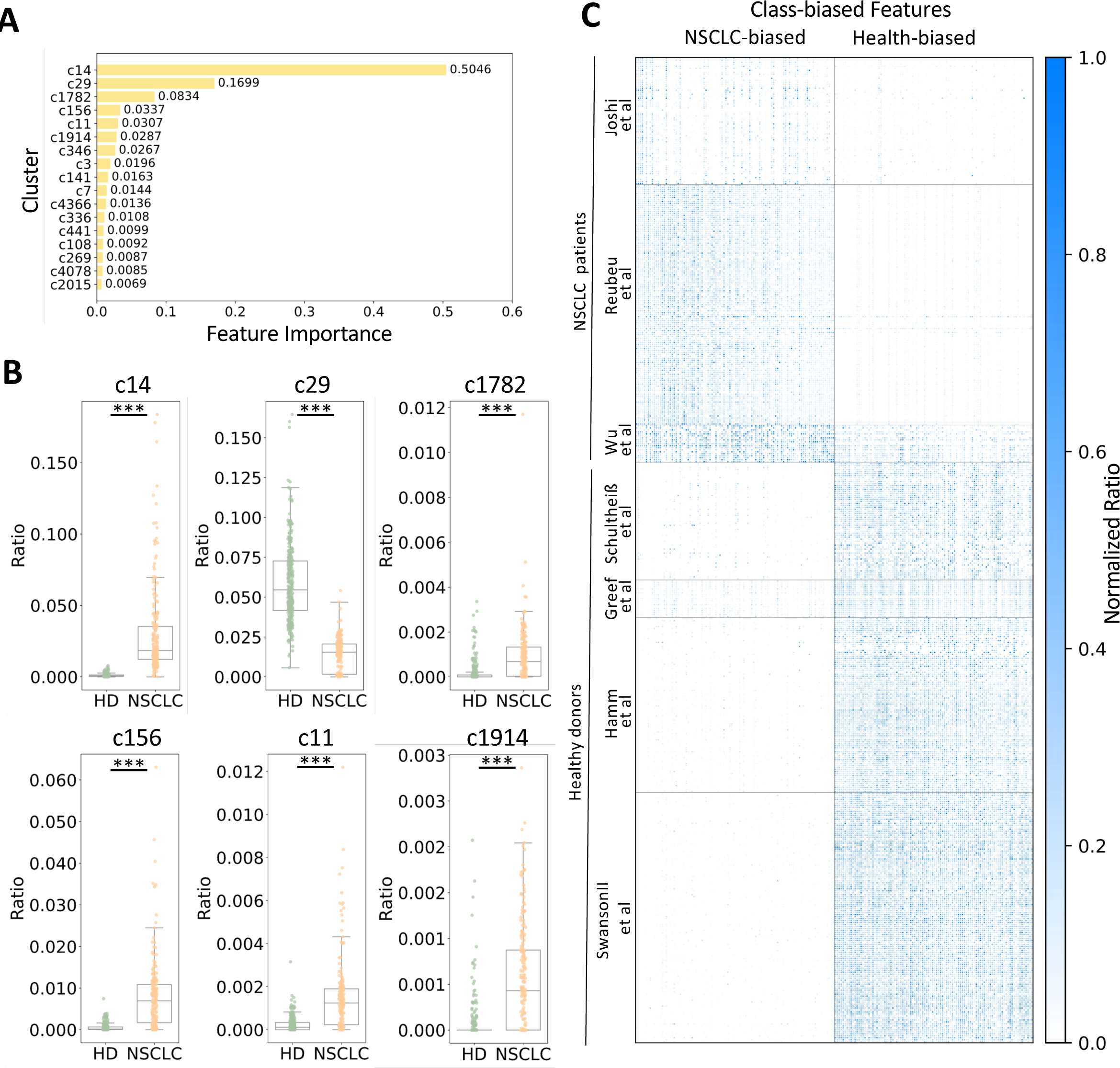
Assessment of cluster features in NSCLC patient classification. **A**, Feature importance. The Y-axis shows the importance of the XGBoost features in descending order. **B**, The six most important features. Each dot represents the proportions of healthy (HD) and disease (NSCLC) donors exhibiting those features. A t test was performed to compare the proportions between the HD and NSCLC cohorts (\*\*\**p*≤0.001). **C**, Heatmap of the top 100 significant disease and healthy (HD) cluster features. *p* values were calculated with the t test; by sorting the *p* values in ascending order, the ranks of the corresponding features are determined. The value of each cell is the normalized ratio from 0 (white) to 1 (blue). The left Y-axis shows each donor from the healthy and NSCLC cohorts grouped by the corresponding study.

**Table 1.**
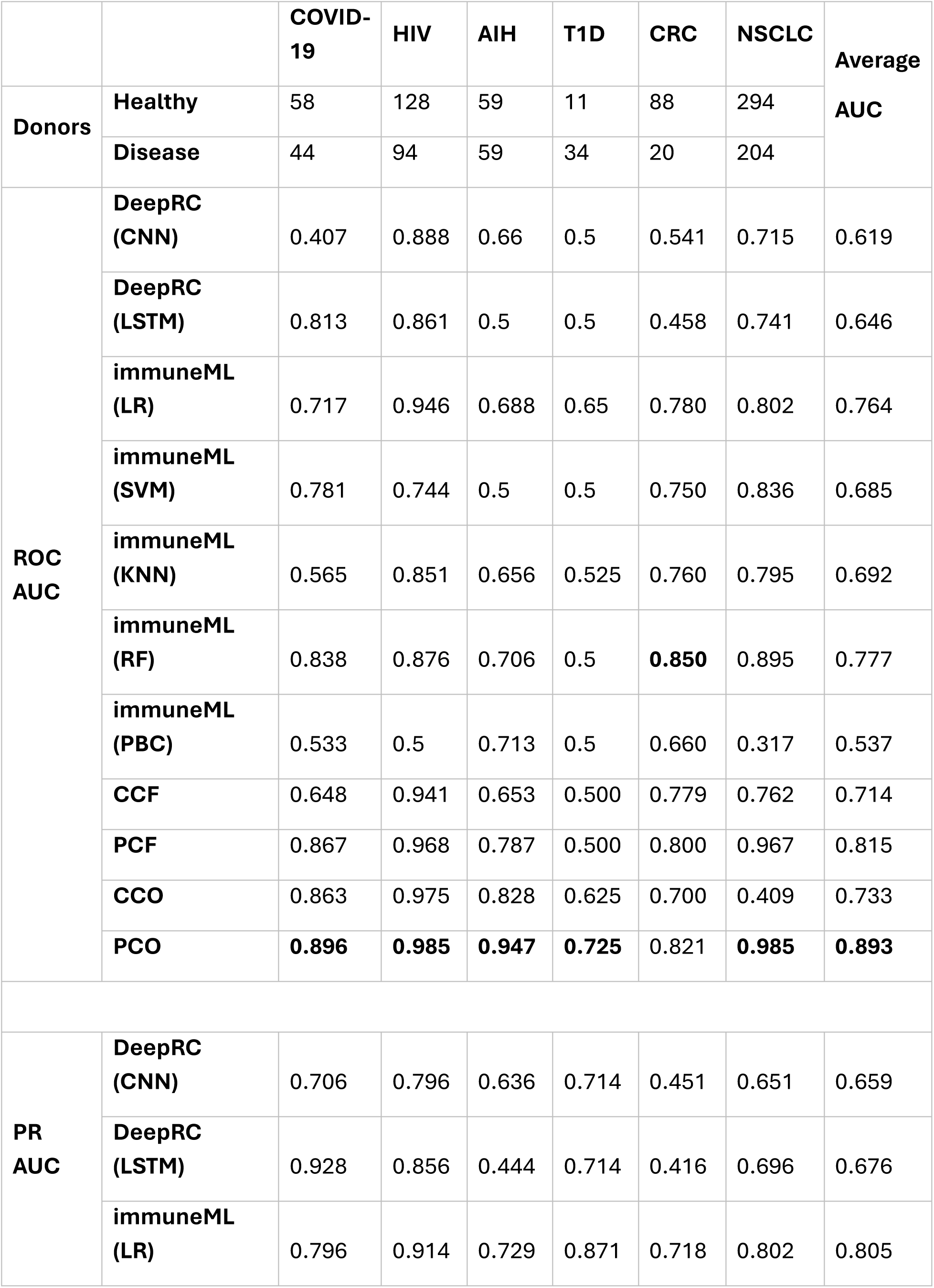
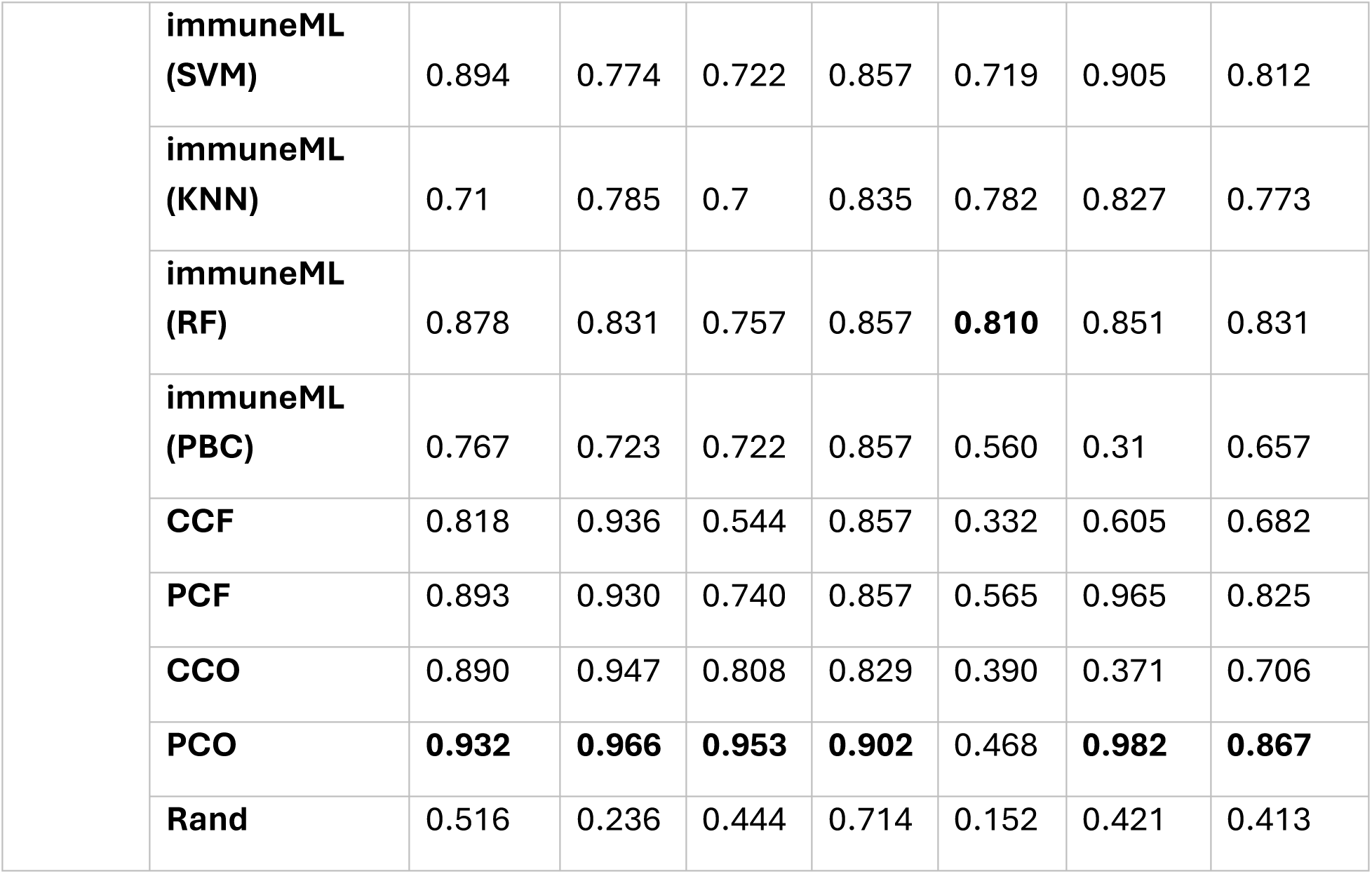
Summary of performance metrics. Performance of previously published methods (DeepRC, immuneML) using various settings, along with the classifiers developed in this study, each trained on one of the four features (CCF, PCF, CCO, and PCO) and applied to six diseases (COVID-19, HIV, AIH, T1D, CRC, and NSCLC). The numbers of donors are listed under each disease. Areas under both the receiver operating characteristic (ROC AUC) and precision-recall (PR AUC) curves are given. For the PR AUC values, the expected performance of a random predictor (Rand) is given by the ratio of positive to negative donors. The abbreviations in the second column are as follows: convolutional neural network (CNN), long short-term memory (LSTM), logistic regression (LR), support vector machine (SVM), K-nearest neighbor (KNN), random forest (RF), and probabilistic binary classifier (PBC).

### Classifiers trained on PCOs were robust against batch eDects

Because our large NSCLC dataset was composed of data from multiple studies, it provided the opportunity to systematically explore the robustness of the different features against batch effects: effects that arise from differences between samples that are not rooted in the experimental design and can have various sources^43^. To this end, we sampled all possible splits of the datasets where the test set consisted of the data from one healthy study and one NSCLC study, while the data from the remaining studies were used for training. This process resulted in twelve splits with various numbers of AIRs in the training and test sets. These results indicated that, in terms of the ROC (**Fig. 3E**) and PR AUCs (**Fig. 3F**), the classifiers based on PCF and PCO features performed well overall, but those based on the CCF and CCO features performed inconsistently, in agreement with the data shown in **Figs. 3C-D**. Furthermore, two of the twelve splits showed that it was possible to combine the NSCLC studies in such a way that the PCO model failed to generalize to the test data. Interestingly, in these two splits, the size of the training dataset was much smaller than that of the test dataset, which was consistent with the findings for the other diseases like T1D and CRC, again emphasizing the need for sufficient training data. Overall, however, the performance of the PCO-based classifier was more robust than that of the alternative new classifiers.

### Cancer data reveals a surprising relationship between healthy-biased clusters and cancer antigen specificity

The underlying AIRs can be investigated using the XGBoost importance to identify clusters of interest. Because each feature corresponds to an AIR cluster, we hypothesized that AIRs within clusters corresponding to important features with high importance values might be more likely to target antigens that are specific to the disease in question. To test this hypothesis, we again examined the NSCLC dataset, for which we had the largest amount of data. First, we ranked the PCO clusters by their feature importance (**Fig. 4A**) and examined the proportions of healthy- and NSCLC-derived TCRs in each. We observed clusters both with significantly more TCRs from healthy donors (“healthy-biased” clusters) and with significantly more TCRs from disease donors (“NSCLC-biased” clusters) using a t test p value cutoff of 0.001 (**Fig. 4B**). The features of these class-imbalanced clusters are shown as a heatmap in **Fig. 4C**. Next, using these class-imbalanced clusters, we searched two databases (McPAS^44^ and VDJdb^45^), whose TCRs are annotated by their targeted antigen and associated disease. Interestingly, we identified strong hits to cancer-associated antigens using the TCRs from both the healthy- and NSCLC-biased clusters, but surprisingly, there were more hits from the healthy-biased clusters than from the NSCLC-biased clusters (**Fig. 5A)**. **Fig. 5B** illustrates some of these hits, which included antigens such as MLANA^46^, EphA2^47^, BST2^48^, TKT^49^ and IGF2BP2^50^, which have been reported as prognostic markers for NSCLC. These findings demonstrate the further application of PCO-based features as a means of identifying potential cancer-fighting T cells.

**Figure 5.**
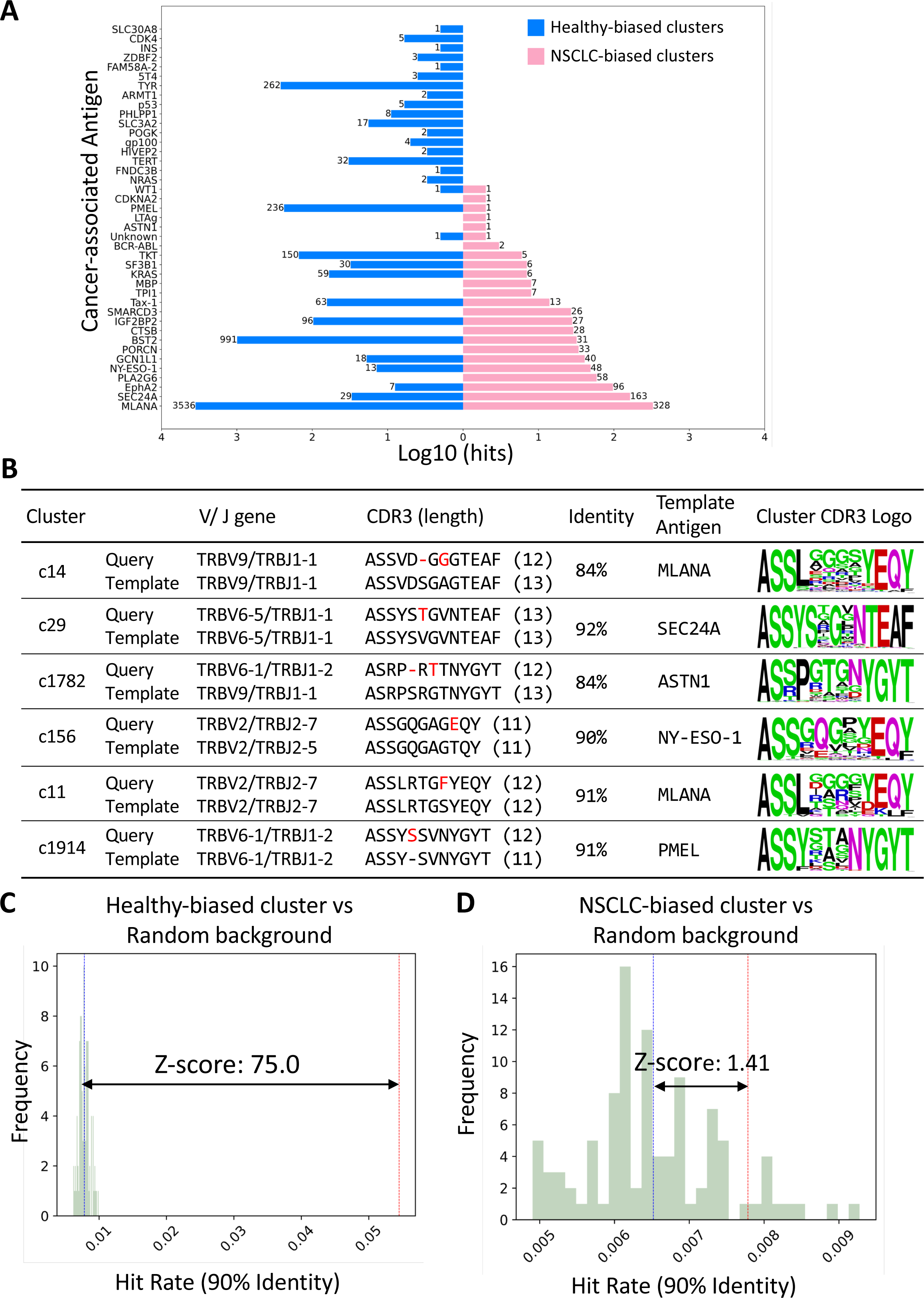
Functional annotation of healthy- and NSCLC-biased clusters. **A**, Histogram of hits to TCRs targeting the indicated antigens from healthy- (blue bars) and NSCLC-biased (pink bars) clusters after querying the McPAS and VDJdb cancer databases. Each count represents one database hit. **B**, Sequences from important clusters matching known cancer-targeting antigens, as shown by their alignments and sequence logos based on all members of the same cluster. **C**, Hit rates of TCRs from healthy-biased clusters to cancer-targeting TCRs (vertical red line) and the corresponding hit rates of 100 randomly selected sets of queries from the same donors (green bars with blue vertical lines representing the means). The horizontal arrow indicates the Z score. **D,** Hit rates of TCRs from NSCLC-biased clusters to cancer-targeting TCRs (vertical red line) and the corresponding hit rates of 100 randomly selected sets of queries from the same donors (green bars with blue vertical lines representing the means). The horizontal arrow indicates the Z score.

We speculated that T cells that target cancer-associated antigens might be more common in healthy donors if, in NSCLC patients, they migrate from the peripheral blood toward tumor-presenting tissues (i.e., the lungs). To test this idea, we repeated the database queries using randomly selected TCRs from healthy donors. Compared to TCRs from healthy biased clusters, randomly selected TCRs from healthy donors resulted in dramatically fewer hits (Z score = 75), as shown in **Fig. 5C**. In contrast, TCRs from NSCLC-biased clusters exhibited a similar level of cancer-associated hits to randomly selected TCRs from donors with NSCLC (Z score = 1.41), as shown in **Fig. 5D**. These observations were not qualitatively sensitive to the similarity threshold used to define a hit (**Fig. S4**). Taken together, they show that the healthy-biased TCRs are indeed distinct from typical healthy donor-derived TCRs, supporting the notion that they would have the potential to migrate out of the periphery in cancer patients.

We next performed an analogous analysis for colorectal cancer (CRC) cases (**Fig. S5-6**). Consistently, TCRs from healthy-biased clusters had significantly more cancer-related hits than did randomly selected TCRs from healthy donors (Z score = 39.5) (**Fig. S6C**), regardless of the similarity threshold (**Fig. S7**). A recent report showed that several cancer-specific TCRs could target multiple tumor types via the HLA A*02:01-restricted epitopes EAAGIGILTV, LLLGIGILVL, and NLSALGIFST from Melan A (MLANA), BST2, and IMP2 (IGF2BP2) and that PBMCs from healthy donors expressed such TCRs^51^. Consistently, many TCRs from healthy-biased, important clusters were predicted to target MLANA and BST2 or MLANA and IGF2BP2 via highly similar epitopes in both the NSCLC (**Fig. S8**) and CRC (**Fig. S9**) datasets. Taken together, these results strongly suggest that the TCRs in the healthy-biased clusters migrate out of the peripheral blood to infiltrate tumors in individuals with NSCLC and CRC. These findings highlight the interpretability of our ML classifiers to identify underlying mechanisms and potentially therapeutic immune cells.

## Discussion

Liquid biopsies that can detect specific diseases from blood have the potential to reshape the future of medical diagnosis. Adaptive immune cells, which circulate in peripheral blood, are highly sensitive to a broad range of diseases. However, harnessing this sensitivity in a reproducible and general manner has presented a challenge, due to the high diversity and low inter-donor sharing of AIRs. Moreover, the natural inclination to gather more extensive and larger datasets for machine learning purposes is at odds with the goal of achieving widespread mutual sharing among all donors. This problem is exacerbated by the use of the clonotype nomenclature, which separates AIRs into discrete clusters defined by their V and J gene usage and CDR3 length. Our results demonstrate that an AIR representation that incorporates the extended networks of paratopes allows donors to be connected and improves classifier performance compared to clonotype-based classifiers.

We found that there was obvious improvement with additional training data, which is encouraging given the rapid growth of publicly available AIR data. Beyond a threshold value of 200 donors, the PCO-based classifiers were very robust, achieving ROC AUC values of 0.985 (HIV and NSCLC). This suggests that initial clinical validation of specific diseases could be performed with smaller numbers of donors (e.g. 20 patients) and then scaled up to larger numbers (e.g. 50 patients) based on initial results.

The network view of adaptive immunity makes sense from the perspective of host defense, as pathogens represent a diverse and unpredictable set of threats. Indeed, Immune Network Theory, originally proposed by Jerne and Hoffmann helped to explain the immune system’s ability to distinguish self from non-self^52,53^. Here, we see from the perspective of disease classification that dense features based on paratope networks generalize much better to new donors than sparse features based on less connected clonotypes and that these observations were consistent across three broad categories of disease.

In addition to these general observations, we found that in the two cancer datasets, clusters that corresponded to important features included both healthy- and disease-biased clusters. Typically, in repertoire analysis, there is a tendency to focus on clusters that are overrepresented in patients^54, 55^. However, we observed the reverse: healthy-biased clusters were important for disease classification as well. One possible explanation for the importance of healthy-biased clusters is that they are not relevant to cancer and are thus downregulated in individuals with cancer. However, the opposite interpretation—that these clusters are relevant to cancer but have migrated out of the peripheral blood (e.g., to the site of the tumor)—is also conceivable. The latter explanation is consistent with the observation that tumor-infiltrating lymphocytes (TILs) have been observed in almost all cancers^56^. Although we cannot determine what happens to the T cells missing from cancer patients, the extremely high hit rate of cancer-associated TCRs in healthy-biased clusters with respect background healthy TCRs supports the idea that these T cells leave the periphery of cancer patients as a result of targeting tumors. Moreover, the close resemblance of the TCRs identified here to those found to be reactive to multiple types of cancer^51^ further strengthens this argument. The notion that tumors can affect global migration from peripheral blood has been demonstrated in brain cancer and metastatic melanoma^57^. In one NSCLC source report^42^, the authors discovered that many expanded intratumoral TCRs were detectable in blood samples at the time of lung tumor resection from NSCLC patient, consistent with the notion that T cells in peripheral blood, and TCR-based classification of cancer cells from PBMCs, can be used for disease monitoring and personalized immunotherapy development.

### XXX

Undoubtedly, there are limitations to this study that suggest future directions. One concerns the focus on specific disease and limited sample sizes. In this study, we focused on classifying single disease patients from a healthy donor cohort. However, future work must also explore the classification of different cancer types or multiple diseases simultaneously. To achieve this, larger and higher-quality AIR datasets with accompanying clinical information, such as tumor stage, treatment history, and survival outcomes, will be necessary. Expanding the scope of the study to include more diverse disease types and larger cohorts will also help validate the generalizability of our findings and provide a more comprehensive understanding of the TCR repertoire’s role in disease diagnosis and monitoring. Another area we have not yet explored is the combination of BCR and TCR information. Our study focused on BCRs for infectious diseases and TCRs for autoimmune diseases and cancer; however, it would be valuable to investigate the potential of combining both BCR and TCR information, including paired (light-heavy, alpha-beta) chains, to train the classifier. By integrating data from both arms of the adaptive immune system, future studies may potentially improve classification performance and provide a more holistic view of the immune response in various disease states. In addition, incorporating more advanced ML model is of interest. With the advancement of large language models, it will be interesting to explore whether paratope networks emerge naturally from training rather than through explicit calculation of the adjacency matrix. By leveraging these advanced models, we may capture more complex patterns and interactions within TCR and BCR repertoires. In parallel, extending our approach through use of deep learning models, such as graph neural networks, may further improve diagnostic performance.

XXX In conclusion, our study represents a proof of concept showing the use of paratope networks improves the robustness of disease classifiers with the potential to diagnose infectious diseases, autoimmune disorders, and cancer from adaptive immune receptor repertoire data. These findings highlight the potential of our method to provide a deeper understanding of the role of the adaptive immune system in three major disease states and contributes to the development of more accurate and widely applicable liquid biopsies. Moreover, since most liquid biopsies utilizing peripheral blood utilize only the serum component, our technology is entirely compatible with such tests and may well add breadth and sensitivity to existing tests. As the field of AIR repertoire analysis continues to evolve, we anticipate that further refinement will enhance the performance and generalizability of this approach, ultimately leading to improved patient care and outcomes.

---

**Figure S1.**
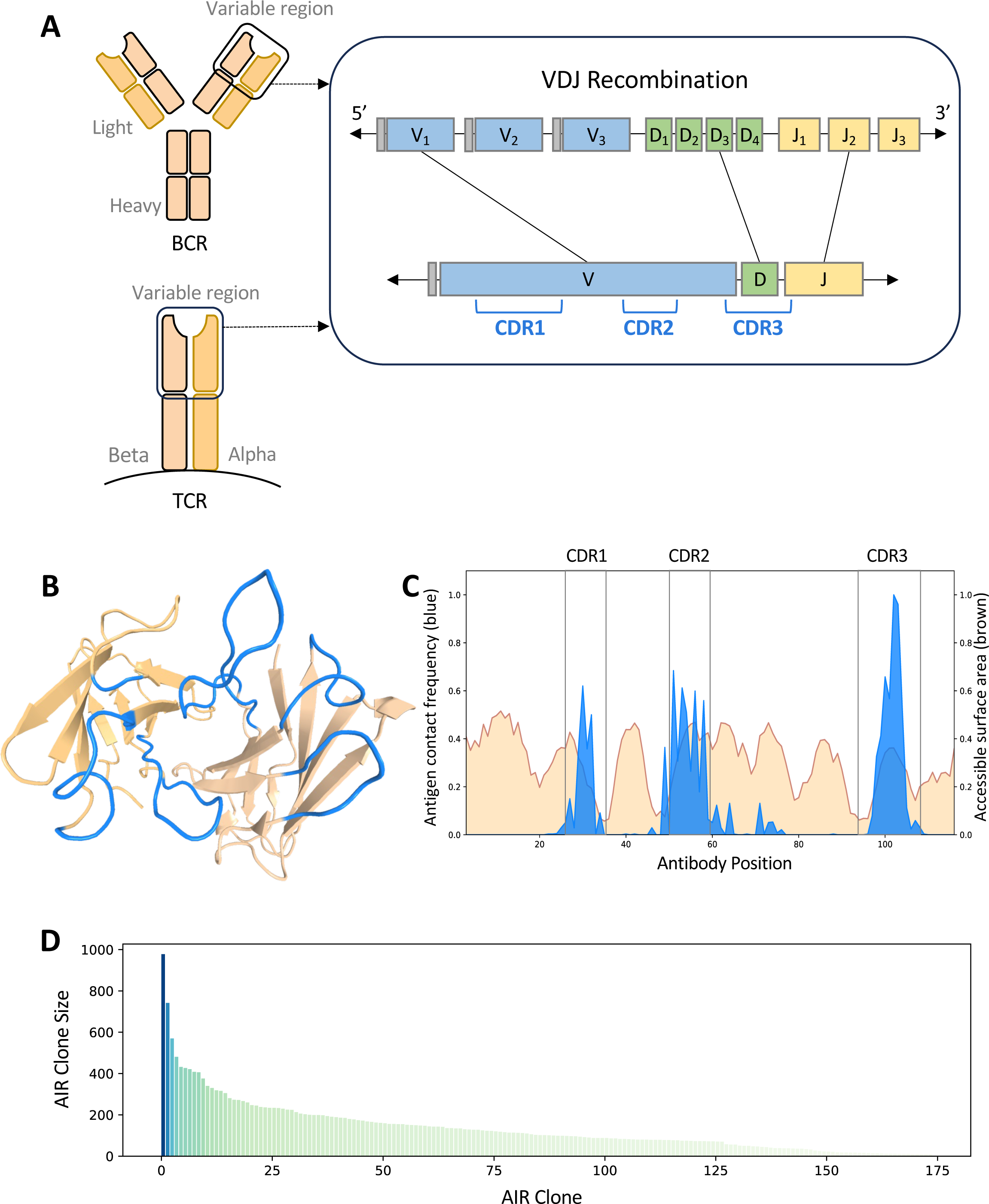
Adaptive immune receptors. **A**, B-cell receptors (also known as antibodies in their soluble form) consist of two heavy and two light chains, while T-cell receptors (TCRs) consist of a single beta and a single alpha chain. BCR and TCR coding sequences are generated by combinatorial rearrangement of V, D and J genes, which results in diverse complementarity determining regions (CDRs 1-3). **B,** CDRs 1-3 are arranged to form a continuous molecular surface called a “paratope” (in blue). **C,** Contact with an antigen (light blue) most often occurs with the CDRs than with the background assessable surface (light brown). **D,** The distribution of BCRs and TCRs for each donor follows a long-tailed distribution, with the large majority of clones having very low frequencies (clones with counts less than 10 are not shown).

**Figure S2.**
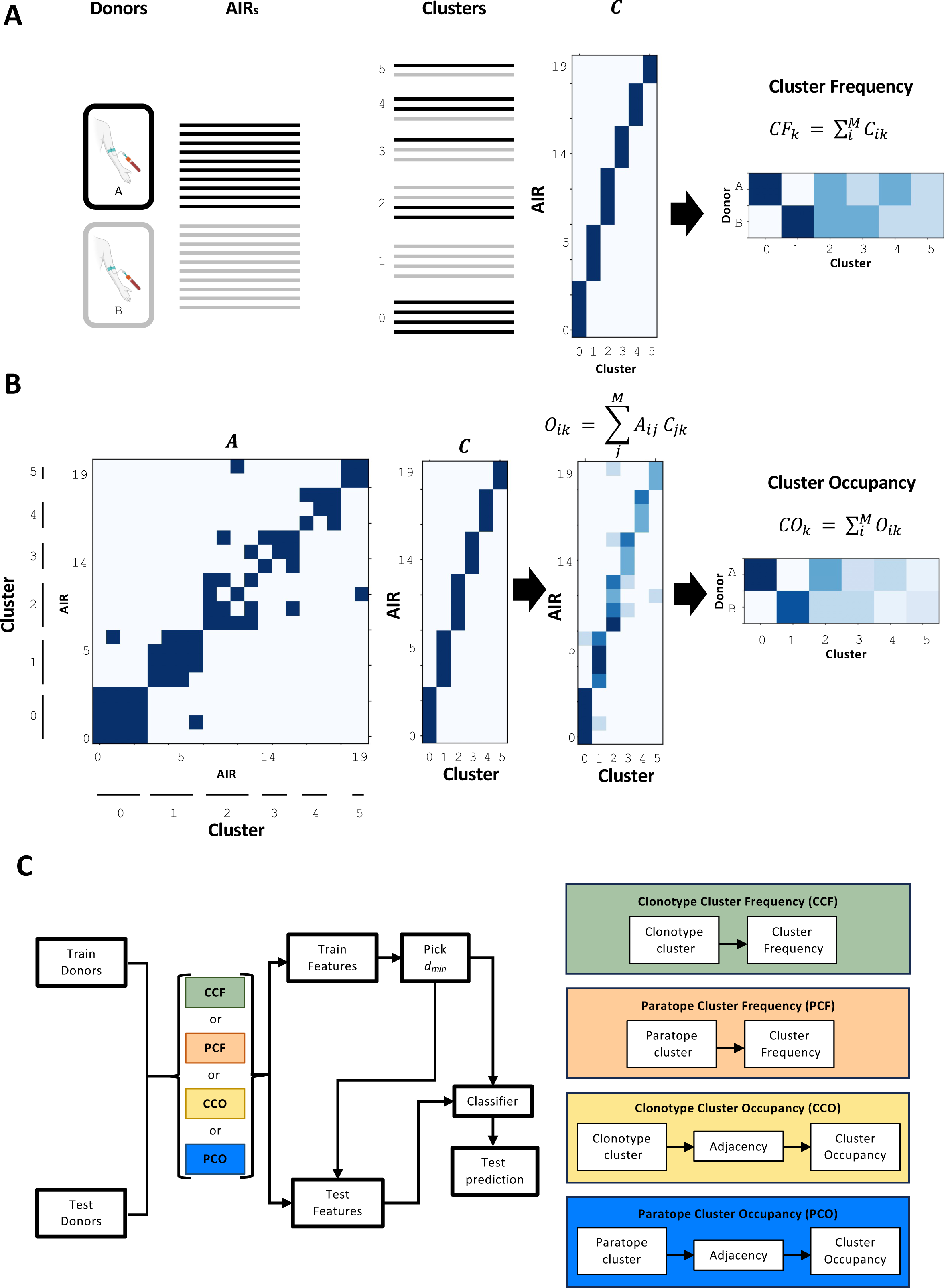
Cluster frequency and occupancy strategy in the classification algorithm. **A**, Cluster frequencies are illustrated with two example donors (A and B) whose AIRs form 6 clusters, which are one-hot encoded in matrix *C*. The cluster frequency feature of a donor is obtained by summing the rows of this matrix that belong to that donor. **B,** Cluster occupancies are derived similarly to cluster frequencies except that we introduce an adjacency matrix describing the pairwise similarities between each AIR. If there are similarities between AIRs belonging to different clusters, the occupancy matrix *O* will differ from *C.* Summing the rows for each donor yields a feature vector that is generally less sparse than that derived from cluster frequencies. **C,** Flowchart of the classification pipeline. Starting from a set of training and test donors, the first step is feature generation using one of four methods, which differ in the type of clustering (clonotype or paratope) and whether the clusters are transformed into features based on frequencies or occupancies. Thereafter, the remaining steps are identical and consist of training/hyperparameter selection and testing the classifier on the test features.

**Figure S3.**
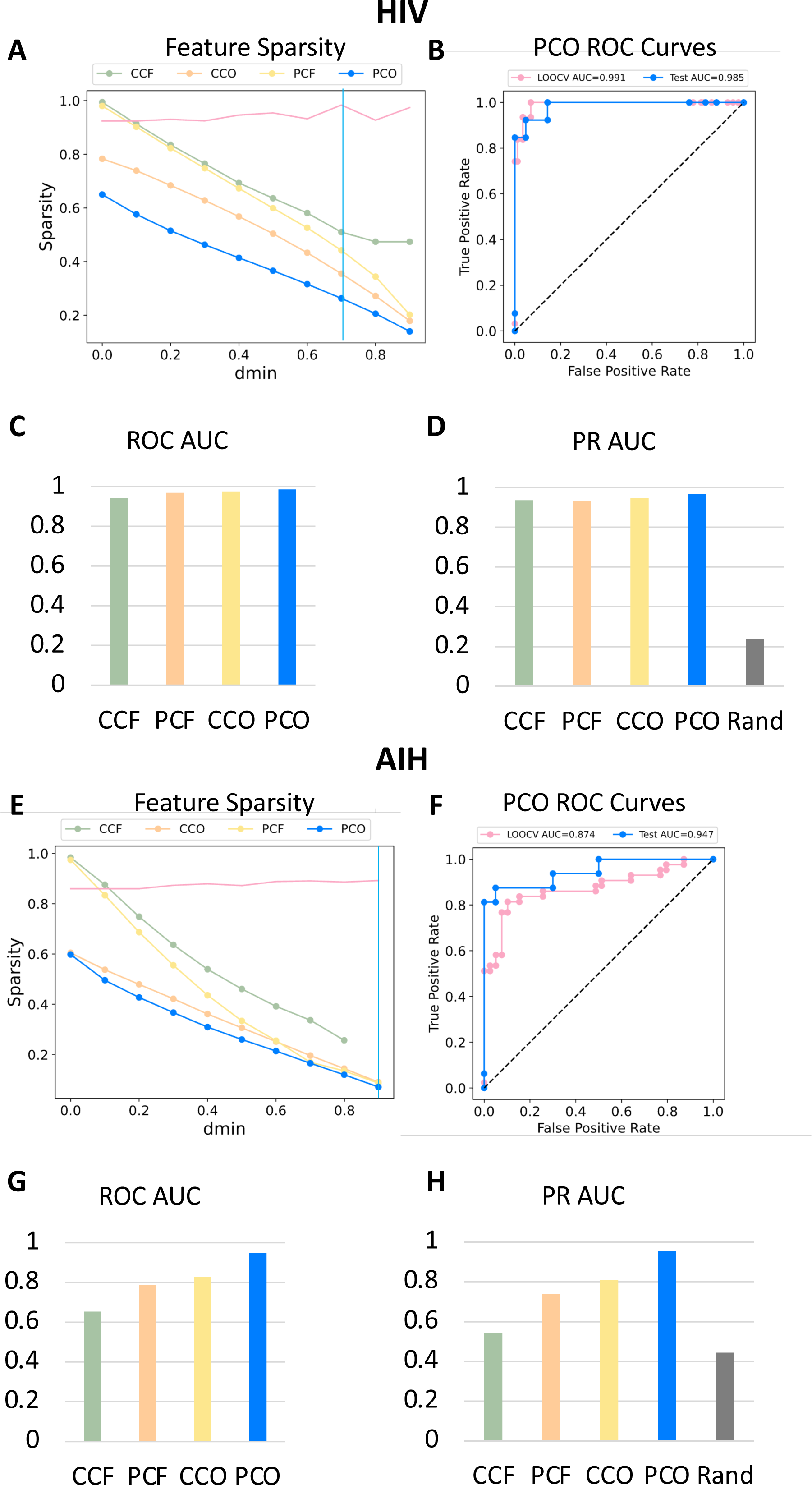
Assessment of HIV diagnosis based on BCRs and AIH diagnosis based on TCRs. **A/E,** The sparsity of each of the four feature matrices as a function of 𝑑_min_, with the vertical cyan line showing the value from LOOCV hyperparameter optimization and the pink line indicating the target function (mean of training ROC AUC and PR AUC). **B/F,** ROC curves for the LOOCV and test predictions using the PCO classifier. **C/G,** ROC AUCs for all four classifiers in the test set. **D/H,** PR AUCs for all four classifiers in the test set; the AUC of a random classifier is indicated as a gray bar. Impact of occupancy on important features.

**Figure S4.**
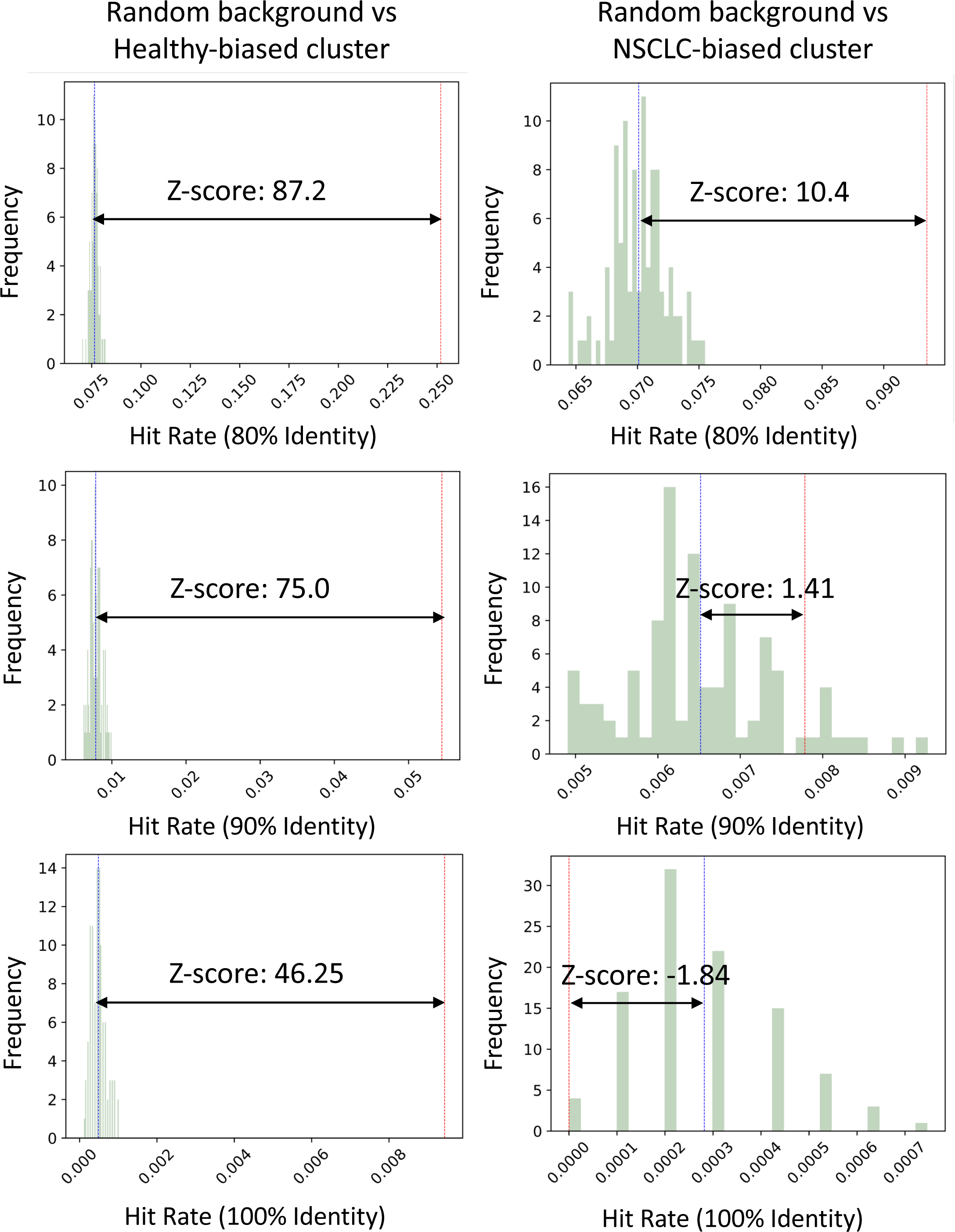
Assessment of T1D and CRC diagnosis based on TCRs. **A/E,** The sparsity of each of the four feature matrices as a function of 𝑑_min_, with the vertical cyan line showing the value from LOOCV hyperparameter optimization and the pink line indicating the target function (mean of training ROC AUC and PR AUC). **B/F,** ROC curves for the LOOCV and test predictions using the PCO classifier. **C/G,** ROC AUCs for all four classifiers in the test set. **D/H,** PR AUCs for all four classifiers in the test set; the AUC of a random classifier is indicated as a gray bar. Impact of occupancy on important features.

**Figure S5.**
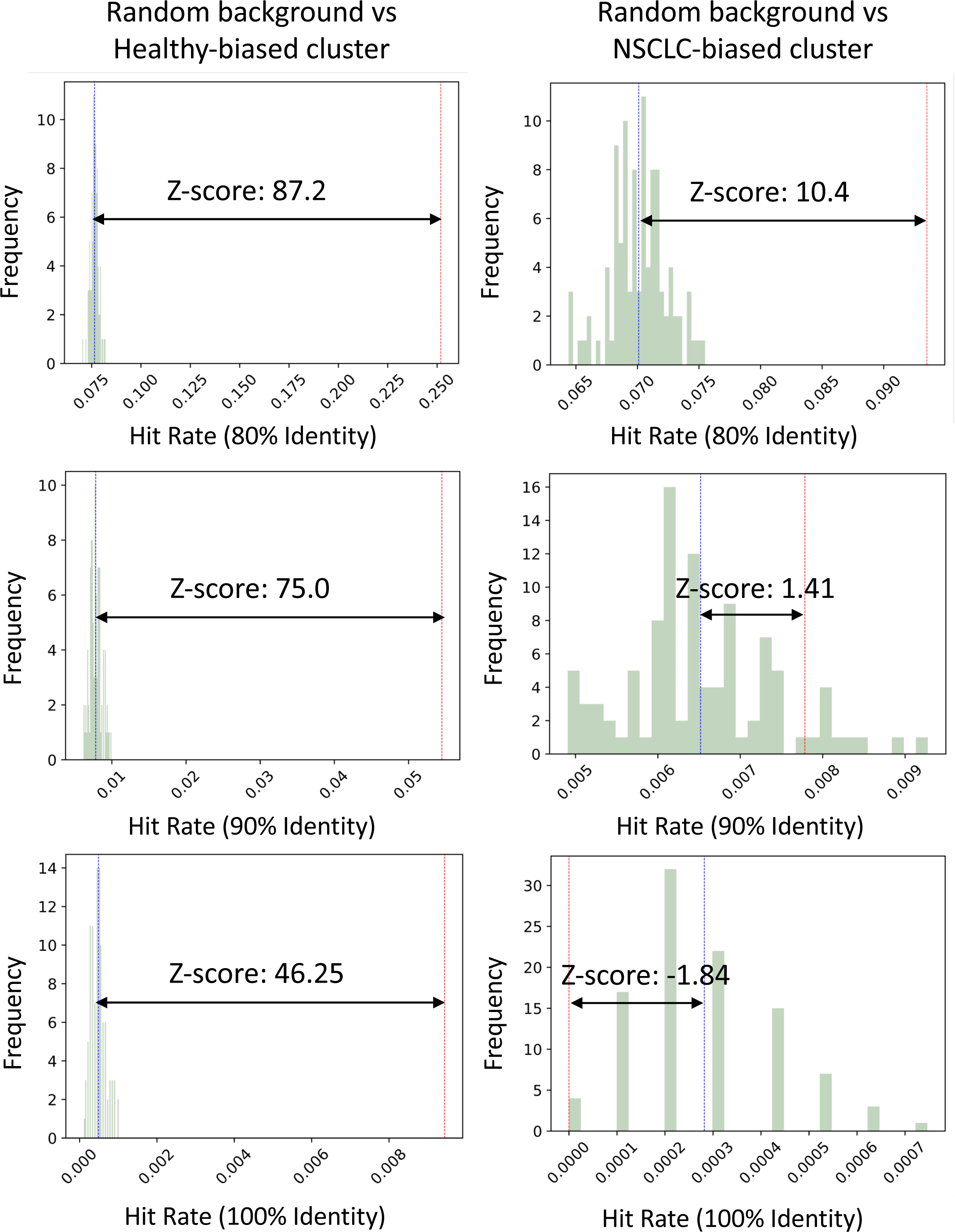
Cancer-associated hit rates from healthy-biased clusters and NSCLC-biased clusters using various similarity thresholds. TCRs from healthy-biased clusters had significantly more cancer-related hits than did those from randomly selected background TCRs under three different CDR3 sequence identity thresholds (80%, 90% and 100%).

**Figure S6.**
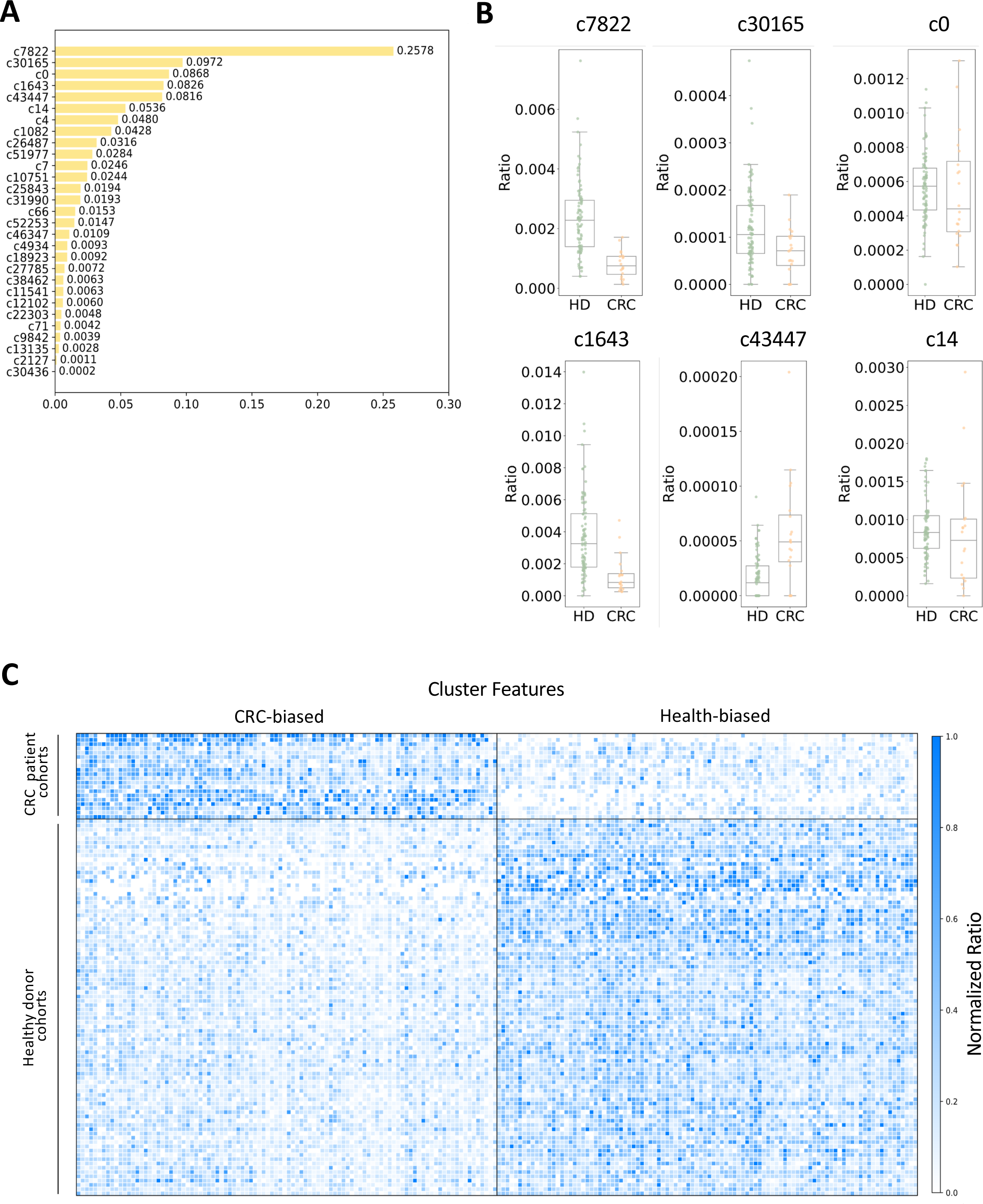
Assessment of cluster features in CRC patient classification. **A**, Feature importance. The Y-axis shows the importance of the XGBoost features in descending order. **B**, The top six most important features. Each dot represents the proportion of healthy (HD) and disease (CRC) donors with that feature. A t test was performed to compare the proportions between the HD and CRC cohorts (\*\*\**p*≤0.001). **C**, Heatmap of the top 100 CRC- and healthy (HD)-biased cluster features. *p* values were calculated with the t test; by sorting the p values in ascending order, the ranks of the corresponding features was determined. The value of each cell is the normalized ratio from 0 (white) to 1 (blue). The left Y-axis shows each donor from the healthy and CRC cohorts grouped by the corresponding study.

**Figure S7.**
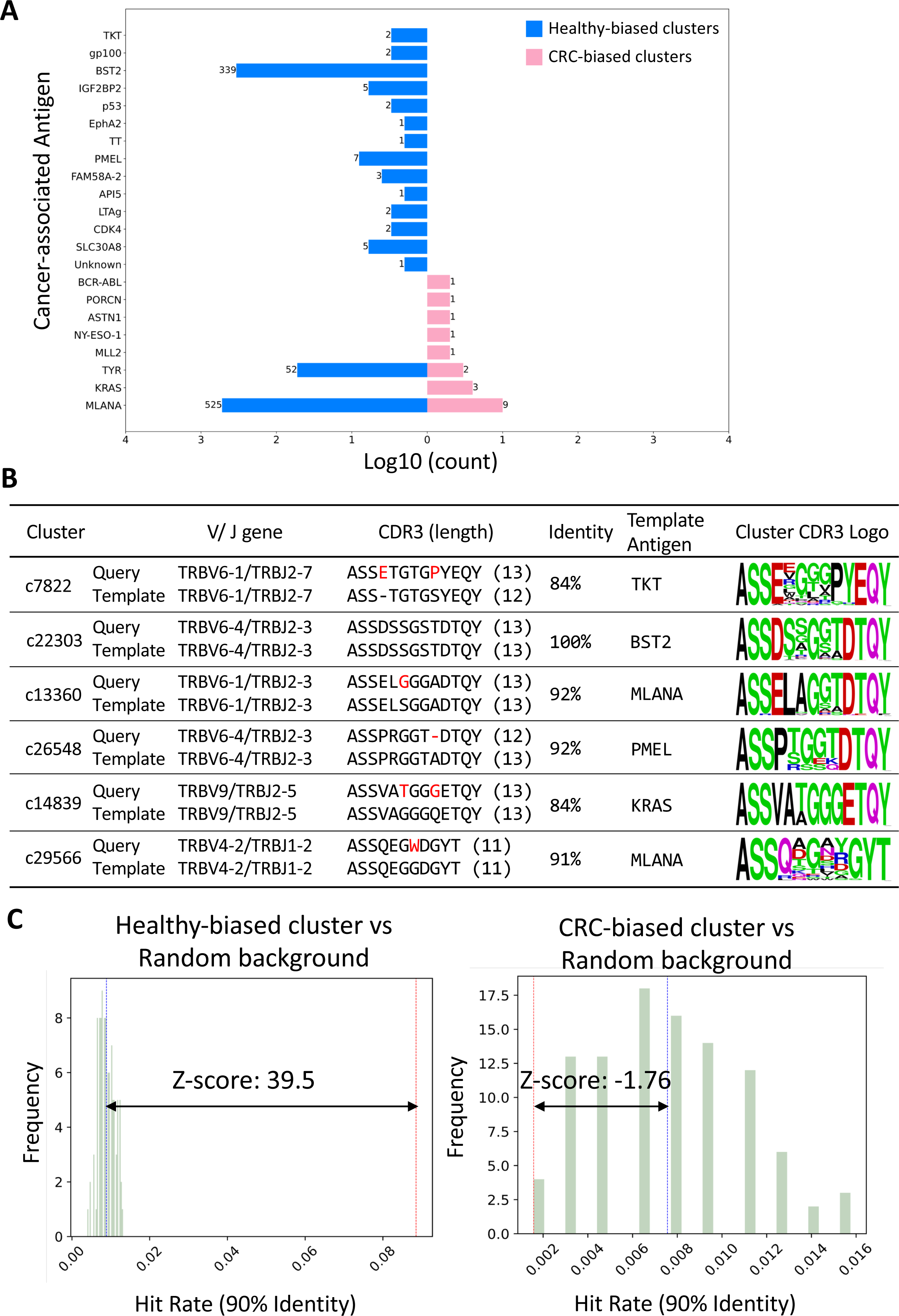
Functional annotation of healthy- and CRC-biased clusters. Histogram of hits to TCRs targeting the indicated antigens from healthy- (blue bars) and CRC-biased (pink bars) clusters after querying the McPAS and VDJdb cancer databases. Each count represents one database hit. **B**, Sequences from important clusters matching known cancer-targeting antigens, as shown by their alignments and sequence logos based on all members of the same cluster. **C**, Hit rates of TCRs from healthy-biased clusters to cancer-targeting TCRs (vertical red line) and the corresponding hit rates of 100 randomly selected sets of queries from the same donors (green bars with blue vertical lines representing the means). The horizontal arrow indicates the Z score. **D,** Hit rates of TCRs from CRC-biased clusters to cancer-targeting TCRs (vertical red line) and the corresponding hit rates of 100 randomly selected sets of queries from the same donors (green bars with blue vertical lines representing the means). The horizontal arrow indicates the Z score.

**Figure S8.**
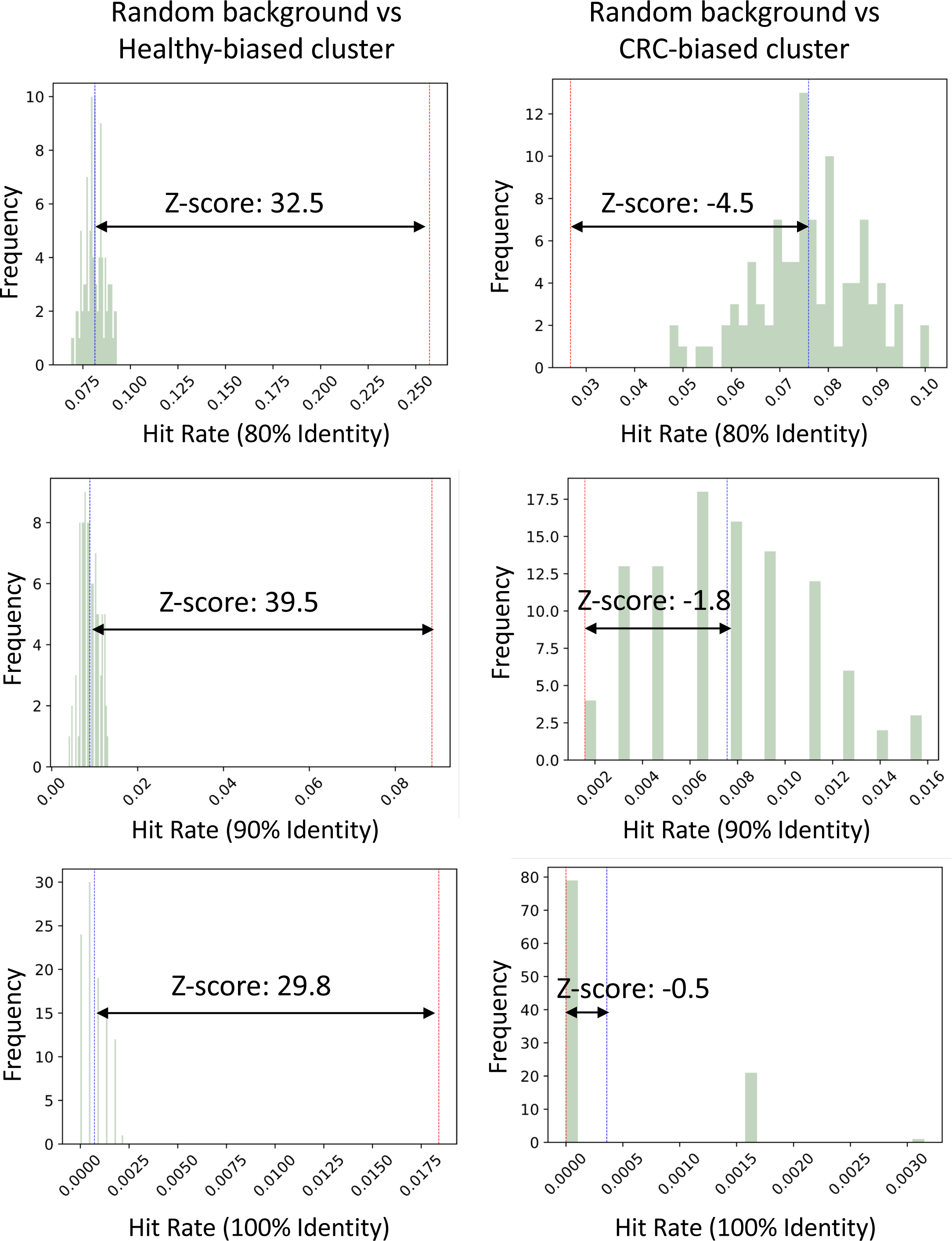
Cancer-associated hit rates from healthy-biased clusters and CRC-biased clusters using various similarity thresholds. TCRs from healthy-biased clusters had significantly more cancer-related hits than did those from randomly selected background TCRs under three different CDR3 sequence identity thresholds (80%, 90% and 100%).

**Figure S9.**
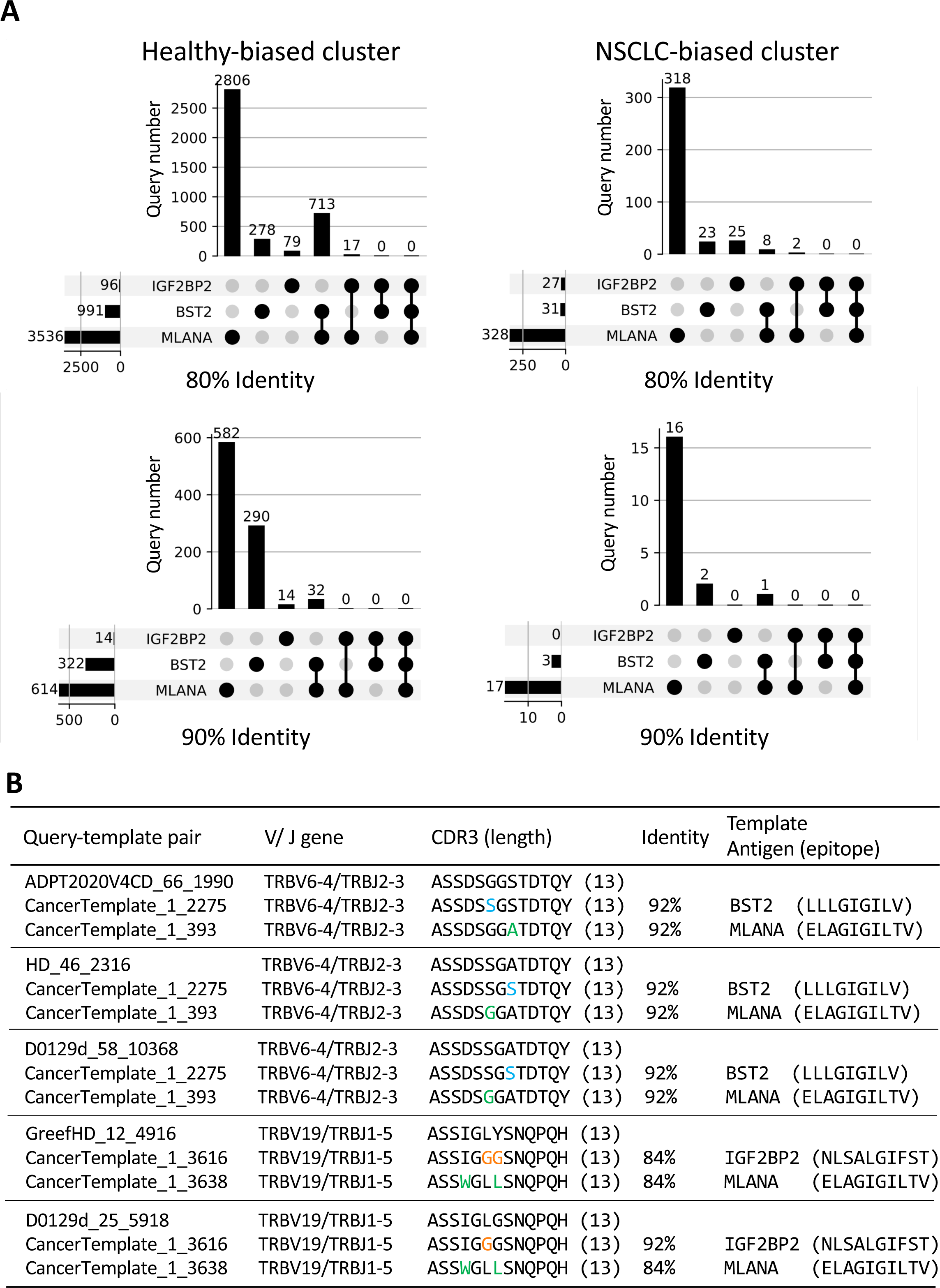
Predicted MLANA, BST2, and IGF2BP2-targeting TCRs. **A**, The UpSet plot shows the TCR hit numbers for the predicted MLANA, BST2, and IGF2BP2-targeting TCRs from healthy (left) and NSCLC-biased (right) clusters under a CDR3 sequence identity of 80 or 90%. Some TCRs are predicted to target two antigens. **B,** Sequence alignments of query-template hits indicating that multiple healthy donors harbor TCRs predicted to target two cancer-related antigens.

**Figure S10.**
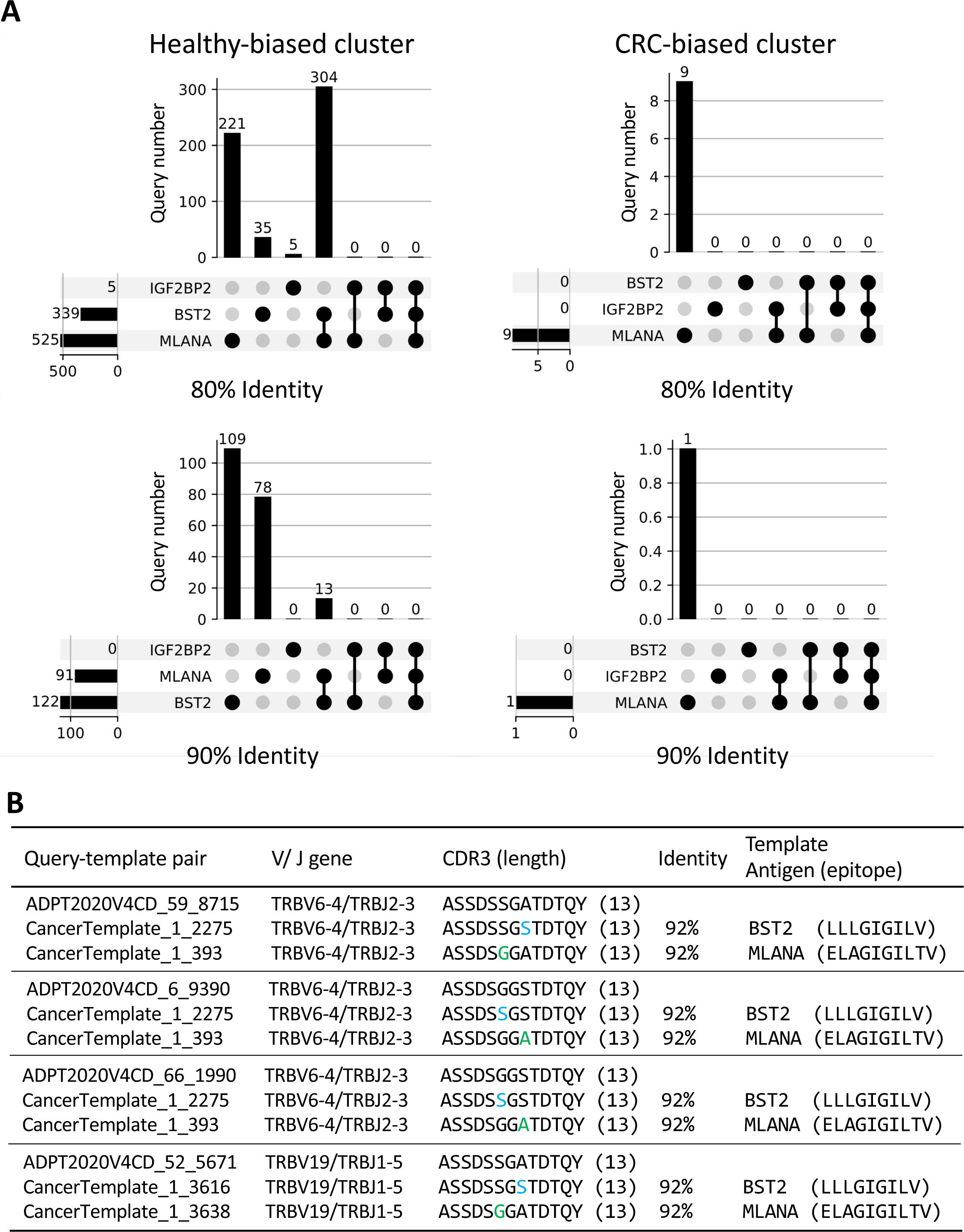
Predicted MLANA, BST2, and IGF2BP2-targeting TCRs. **A,** The UpSet plot shows the TCR hit numbers for the predicted MLANA, BST2, and IGF2BP2-targeting TCRs from healthy (left) and CRC-biased (right) clusters with CDR3 sequence identities of 80 or 90%. Some TCRs are predicted to target two antigens. **B,** Sequence alignments of query-template hits indicating that multiple healthy donors harbor TCRs predicted to target two cancer-related antigens.

## Notes

### Competing Interest Statement

DMS has filed patent applications for the clustering and disease classification technologies.

### Summary of Updates

we have undertaken a comprehensive revision to refine our argument and better articulate the innovative nature of our work. Specifically, we have re-framed the aim of study as a novel liquid biopsy, improved the description of the novel approach in more accessible terms, and provided a illustration based on new data to highlight the differences to the conventional (clonotype) approach.

