## Supplementary Material for "Robust diagnosis of infectious disease, autoimmunity and cancer from the paratope networks of adaptive immune receptors"

### Table S1. AIR datasets

| Disease | Class | Donors | AIRs | Chain | Bioproject | References |
|---|---|---|---|---|---|---|
| **Coronavirus disease 2019 (COVID-19)** | Disease | 44 | 106,640 | BCRH | JGAS000593 | 1 |
| | Healthy | 58 | 174,139 | | | |
| | Total | 102 | 280,779 | | | |
| **Human immunodeficiency virus (HIV)** | Disease | 94 | 5,202,133 | BCRH | PRJNA486667 | 2 |
| | Healthy | 128 | 7,934,405 | | | |
| | Total | 222 | 13,136,538 | | | |
| **Autoimmune hepatitis (AIH)** | Disease | 59 | 338,541 | TCRB | PRJEB37143 | 3 |
| | Healthy | 59 | 390,507 | | | |
| | Total | 118 | 729,048 | | | |
| **Type 1 Diabetes (T1D)** | Disease | 34 | 2,161 | TCRB | PRJNA758093 | 4 |
| | Healthy | 11 | 408 | | | |
| | Total | 45 | 2,569 | | | |
| **Non-small cell lung cancer (NSCLC)** | Disease | 204 | 6,734,867 | TCRB | https://doi.org/10.21417/AR2019NC<br>PRJNA544699<br>PRJNA477518<br>PRJEB37143<br>PRJNA767497<br>https://doi.org/10.21417/ADPT2020V4CD<br>http://doi.org/10.21417/PAS2021STM | 3,5-10 |
| | Healthy | 294 | 4,120,597 | | | |
| | Total | 498 | 10,855,464 | | | |
| **Colorectal cancer (CRC)** | Disease | 20 | 137,622 | TCRB | https://doi.org/10.21417/ADPT2020V4CD | 10 |
| | Healthy | 88 | 1,100,012 | | | |
| | Total | 108 | 1,237,634 | | | |

***Supplementary text***

### Classification of HIV patients and healthy donors from BCR data

HIV is a retrovirus that infects helper T cells, resulting in a chronic infection that, if untreated, results in acquired immunodeficiency syndrome (AIDS). Broadly neutralizing antibodies against HIV often have distinct paratope features, including long CDRH3 regions[1]. Here, we utilized a large-scale study involving BCR heavy chain sequences from the PBMCs of 94 HIV+ (5,202,133

BCRs) and 128 healthy uninfected (7,934,405 BCRs) donors[2]. After random splitting, 155 donors were used for training, and 67 were held out for testing. Compared with the COVID-19 dataset, the HIV dataset is very large, not only in terms of the number of donors but also in terms of the sequencing depth (roughly twenty times that of the COVID-19 study). This presented a problem for the computation of the adjacency matrix, which has a quadratic time complexity with respect to the number of AIRs. To circumvent this problem, we used cluster downsizing as a preprocessing step. Here, we set $d_{min}$ = 0.05, which reduced the number of AIRs from 13,136,538 to 4,424,328. Larger values of $d_{min}$ would have reduced the number of AIRs further while also removing all AIRs for one or more of the training donors. The reduced HIV dataset was used in all subsequent calculations. **Fig. S3A** shows that the feature sparsity was qualitatively lower for CCO and PCO than for CCF or PCF, implying that the occupancy step increases the donor diversity of the clusters, effectively connecting clusters that would otherwise be less diverse, regardless of whether clustering is performed according to clonotypes or paratopes. As with the COVID-19 assessment, the LOOCV target function reached a maximum at a $d_{min}$ value of 0.7. **Fig. S3B** shows that the PCO-based classifier achieved a nearly perfect ROC AUC of 0.985 in the test donor AIRs and that the training LOOCV and test ROC curves overlapped well, indicating that the classifier was not overfitted. As shown in **Figs. S3C-D**, all four models performed similarly in terms of both the ROC AUC and the PR AUC.

**Classification of Autoimmune hepatitis patients and healthy donors from TCR data**

Autoimmune hepatitis (AIH) is a chronic liver disease that is thought to be driven by autoreactive T cells[3]. Here, we utilized a study involving TCR beta chains from the PBMCs of 59 AIH patients (338,541 TCRs) and 59 healthy donors (390,507 TCRs)[4]. After random splitting, 82 donors were used for training, and 36 were held out for testing. **Fig. S3E** shows that the sparsity of the features at low donor diversity thresholds ($d_{min}$) separated into two groups (CCF/PCF and CCO/PCO); however, at larger values, all but the CCF-based feature sparsity values converge to a single value below 0.1%, while the LOOCV target function gradually increased. **Fig. S3F** shows that the PCO classifier achieved a ROC AUC of 0.947 on test donors and even outperformed the training LOOCV.

**Fig. S3G-H** shows that the ROC and PR AUCs for the PCO-based model were superior to those of the CCF, PCF, and CCO models.

**Classification of Type 1 diabetes and healthy donors from TCR data**

Type 1 diabetes (T1D) is an autoimmune disease in which T cells target insulin-producing cells in the pancreas[5]. Unlike the datasets used in the other assessments, which employed unsorted PBMCs, the T cells used here were enriched by culturing with human islet antigen. A total of 2,161 TCRs and 408 TCRs sequenced at the single-cell level from 34 T1D patients and 11 healthy controls, respectively, were used[6]. After splitting the data randomly, 31 donors were used for training, and 14 were used for testing. As was observed for the COVID-19 experiment, the low sequencing depth here resulted in highly sparse clonotype clusters and paratope clusters, populated mostly by single donors. However, the occupancy step (CCO, PCO) produced features with greater donor diversity and reduced sparsity (**Fig. S4A**). The performance on test data was lower for all models than for the other diseases discussed above; even the PCO-based model only reached a ROC AUC of 0.725 (**Fig. S4B**). However, this was significantly better than the ROC AUCs of the CCF and PCF classifiers, which indicated in performance no better than random guessing, while the PR AUCs of the cluster-based classifiers were intermediate between those of the PCO model and a random predictor (**Figs. S4C-D**). Taken together, the results suggest that these T1D data represent the lower limit with respect to training data for acceptable classifier performance but nevertheless suggest that PCOs provide more robust features than CCFs.

**Classification of Colorectal cancer and healthy donors from TCR data**

Colorectal cancer (CRC) is a cancer of the large intestine that is typically diagnosed by tissue biopsy during a colonoscopy or sigmoidoscopy[7]. T cells are known to infiltrate CRC tumors; however, the extent to which these T cells circulate in the blood is poorly understood. Indeed, if disease-specific T cells can be detected in the blood, a TCR-based diagnosis would represent a minimally invasive alternative to existing methods. Here, we utilized TCR beta chain data from a study of 20 CRC patients (137,622 TCRs) and 88 healthy donors (1,100,012 TCRs)[8]. After randomly splitting the data as described above, the TCRs of 75 donors were chosen for training, while those of 33 participants were selected for testing. **Fig. S4E** shows the sparsity of the resulting features, indicating that the PCO and CCO features were less sparse than the other features but that the differences decreased with increasing values of $d_{min}$, similar to the trends observed for the other diseases in this study. The LOOCV target function improved with increasing $d_{min}$ and reached a maximum value at 0.9, which was then selected for constructing the model. **Fig. S4F** shows that the ROC curves for the LOOCV and test sets agree qualitatively and that the test ROC AUC achieved a value of 0.82. When we examined the test ROC AUCs for all four classifiers, the PCO features performed better than the other features (**Fig. S4G**); however, according to the PR AUCs, the PCF classifier performed better than the other three (**Fig. S4H**). This discrepancy between the ROC and PR AUCs is likely derived from the large class imbalance as well as the relatively small size of the CRC dataset.
